# Wnt-dependent spatiotemporal reprogramming of bone marrow niches drives fibrosis

**DOI:** 10.1101/2025.02.12.637594

**Authors:** Bella Banjanin, James Nagai, YeVin Mun, Stijn Fuchs, Inge Snoeren, Joachim Boers, Mayra L. Ruiz Tejada Segura, Hector Tejeda Mora, Anna Katharina Galyga, Adam Benabid, Rita Sarkis, Olaia Naveiras, Marta Rizk, Michael Wolf, Rogerio B. Craveiro, Fabian Peisker, Ursula Stalmann, Jessica E. Pritchard, Hosuk Ryou, Nasullah Khalid Alham, Marek Weiler, Fabian Kiessling, Twan Lammers, Anna Rita Migliaccio, Kishor Kumar Sivaraj, Ralf H. Adams, Eric Bindels, Joost Gribnau, Daniel Royston, Hélène F.E Gleitz, Rafael Kramann, César Nombela-Arrieta, Ivan G. Costa, Rebekka K. Schneider

**Author notes:** contributed equally (Bella Banjanin, James Nagai, YeVin Mun). contributed equally (Hélène F.E Gleitz, Rafael Kramann, César Nombela-Arrieta, Ivan G. Costa, Rebekka K. Schneider).

## Abstract

Bone marrow fibrosis is the most extensive matrix remodeling of the microenvironment and can include *de novo* formation of bone (osteosclerosis). Spatiotemporal information on the contribution of distinct bone marrow niche populations to this process is incomplete. We demonstrate that fibrosis-inducing hematopoietic cells cause profibrotic reprogramming of perivascular CXCL12 abundant reticular (CAR) progenitor cells resulting in loss of their hematopoiesis-support and upregulation of osteogenic and pro-apoptotic programs. In turn, peritrabecular osteolineage cells (OLCs) are activated in an injury-specific, Wnt-dependent manner, comparable to skeletal repair. OLCs fuel bone marrow fibrosis through their expansion and skewed differentiation, resulting in osteosclerosis and expansion of Ly6a+ fibroblasts. NCAM1 expression marks peritrabecular OLCs and their expansion into the central marrow is specific for fibrosis in mice and patients. Peritrabecular stromal b-catenin expression is linked to fibrosis in patients and inhibition of Wnt signaling reduces bone marrow fibrosis and osteosclerosis, possibly being a clinically relevant therapeutic target.

## Introduction

Organ fibrosis contributes to as much as 45% of mortality worldwide and is characterized by extensive tissue remodeling and loss of cellular function (1,2). The cellular origin of extracellular matrix (ECM)-producing cells in various fibrotic diseases has only recently been elucidated through the use of genetic fate-tracing, time-course single-cell RNA- and ATAC-sequencing experiments (3,4,5), which highlight the active tissue remodeling that co-occurs during fibrotic transformation. Bone marrow (BM) fibrosis occurs in a wide spectrum of benign and hematological malignancies, with the myeloproliferative neoplasm (MPN) primary myelofibrosis (PMF) being a prototypical example. In the context of PMF, clonal proliferation of mutated hematopoietic stem cells (HSCs) results in a sustained inflammatory milieu and the activation of BM stromal cells to produce ECM (6–8). Alongside extensive scarring, late-stage disease features *de novo* formation of bone (osteosclerosis), highlighting the plasticity of broadly termed stromal cells within the BM. Yet, the pathogenesis of osteosclerosis is largely unknown. At least two anatomically distinct niches have been described in the BM: the central niche located in the inner BM and the endosteal niche, in close proximity to the bone surface (9). Functional differences between these niches have been proposed (10), raising the possibility that cellular response to injury, e.g. insults occurring in BM fibrosis, differs between anatomically distinct sites.

Here, we provide high resolution single cell and spatial analysis of the BM and specifically ask how stromal subsets change in the presence of fibrosis-inducing hematopoietic cells. For this, we utilize three distinct lineage-tracing stromal Cre-reporters. Previously, we have shown that perivascular Gli1-lineage stromal cells are expanded in kidney, lung, liver, heart, and BM fibrosis (4,5,11), marking a therapeutically attractive cellular target. Additionally, endosteal Gli1-lineage cells were shown to be a major source of osteoblasts in the adult murine skeleton (12). Solid organ fibrosis has traditionally been attributed to the expansion of platelet-derived growth factor receptor-β (Pdgfrb)-positive mesenchymal cells; therefore, we selected PDGFRb as a comprehensive marker for identifying these cells (3,13). Gremlin1-positive (Grem1^+^) cells mark a distinct osteoprogenitor population residing beneath the growth plate, which we hypothesized might be involved in osteosclerosis (14). Using the spatiotemporal resolution provided by these stromal Cre-reporters, we provide evidence that perivascular Cxcl12-abundant reticular cells (CARs) are functionally reprogrammed into fibrosis-driving cells in BM fibrosis and decrease in their frequency, leaving the endothelium exposed. This injury leads to the stepwise activation of injury-specific peritrabecular skeletal progenitor cells. Skeletal progenitor cells exit quiescence, expand and are characterized by skewed differentiation into osteoblasts and fibroblasts, resulting in osteosclerosis and expansion of pro-inflammatory Ly6a^+^ fibroblasts. Mechanistically, peritrabecular progenitor cells are activated in a Wnt-dependent manner with upregulation of β-Catenin. Importantly, Wnt inhibition can reduce BM fibrosis and also inhibit osteosclerosis, being a possible new therapeutic target for myelofibrosis.

## Results

### Distinct spatial macro-niches can be captured by combination of stromal Cre-reporters

We sought to map the BM stromal compartment in homeostatic, unperturbed conditions, and following the induction of BM fibrosis after transplanting thrombopoietin-overexpressing (TPO-OE) hematopoietic stem and progenitor cells or respective controls (empty vector, EV) in lethally irradiated mice. In order to gain a comprehensive spatial overview of the stromal populations, we utilized distinct Cre-reporter strains for lineage fate tracing: 1) Pdgfrb-CreERT2;tdTomato (Pdgfrb;tdTom), a pan-mesenchymal marker well-established in solid organ fibrosis(3,15), 2) Gli1-CreERT2;tdTomato (Gli1;tdTom), a perivascular and endosteal stromal cell population with an established functional role in BM fibrosis (5,12) and 3) Grem1-CreERT2;tdTomato (Grem1;tdTom), described to be a osteolineage-stromal cell population (14).

Within bone, two major macroniches are defined as the metaphyseal region, beneath the growth plate, and the diaphyseal region, containing the central marrow in which the majority of hematopoietic tissue and vasculature resides (>90% of BM) (Figure S1B), (9). The endosteal niche of the metaphysis is composed of trabecular bone and the diaphysis of compact bone.

As expected, the Pdgfrb-reporter marked abundant stromal cells after tamoxifen-induced recombination in homeostatic adult mice (Figure 1A). The central, diaphyseal marrow was filled with Pdgfrb;tdTom^+^ interstitial, reticular-shaped cells, as well as cells surrounding arterial vessels (Figure 1A, diaphyseal panel). Within the metaphyseal region (Figure 1A), Pdgfrb;tdTom^+^ cells were identified as both osteo-lineage cells, and mature osteocytes embedded in the bone matrix, highlighting their active contribution to osteogenesis.

**Figure 1.**
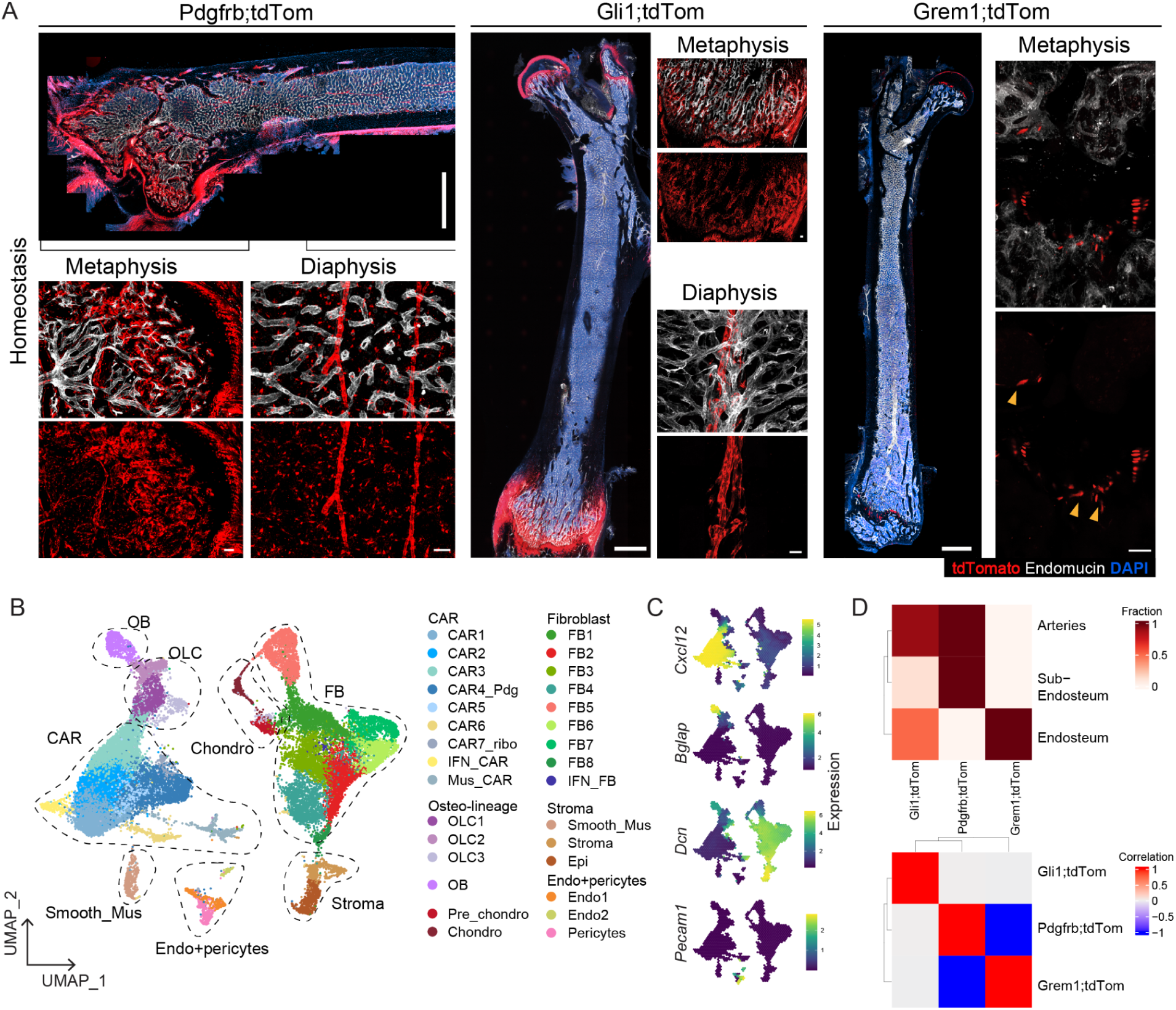
The combination of stromal Cre-reporters provides high granularity of the BM stromal niche. (A) Whole mount imaging of homeostatic (“homeo”) femurs in unperturbed Pdgfr;tdTomato; Gli1;tdTomato and Gremlin1;tdTomato adult mice 21 days after tamoxifen injections. DAPI as nuclear staining (blue) and endomucin (white) to depict vasculature. Yellow arrowheads show sparse Grem;tdTom+ cells emerging from the growth plate. Whole femur overviews scale bar = 1000µm, zoomed-in panels scale bar = 50 µm (B) UMAP representation of 29,742 cells based on the integration of 11 individual scRNAseq libraries (n=40 mice). Major cell type annotations are shown. CAR=CXCL12 abundant reticular cells; FB=fibroblasts; chondro=chondrocytes; epi=epithelial; endo=endothelial; (C) Density of expression values after magic imputation for marker genes of major cell types (D) Deconvolution analysis of scRNAseq data from panel B per distinct Cre-reporter to micro-dissected bulk-RNAsequencing of BM region signatures defined by Baccin et al.(16) Top panel shows the proportion of cells within whole Cre-reporter assigned to each region, bottom panel the correlations between the proportion of Cre-reporters

Gli1;tdTom^+^ cells were residing in the peritrabecular region of the metaphysis (Figure 1A), particularly present in the growth plate and within the compact bone (diaphysis), in line with their known role in postnatal bone formation (12). Gli1;tdTom^+^ cells also had a very specific perivascular position in the diaphysis, wrapping around the central artery; a location previously not appreciated at this resolution. Grem1;tdTom^+^ cells were the least abundant cell population representing singular chondrocyte columns in the growth plate, and a few reticular-shaped cells growing out from the growth plate (Figure 1A), confirming previous reports (14). Importantly, Grem1-tdTom^+^ cells were not detected in the central marrow of mice (Figure 1A, femur overview). The distinct location of these stromal subsets throughout the metaphysis and diaphysis suggested different functions of the targeted stromal cells.

To obtain high-resolution data of these lineage-traced stromal cells in homeostasis and to compare the cellular and transcriptional changes occurring during extensive remodeling of the stroma in BM fibrosis, we performed single-cell RNA sequencing (scRNAseq) on tdTomato^+^ cells isolated from the three Cre-reporters (Pdgfrb;tdTom, Gli1;tdTom and Grem1;tdTom) in homeostasis, BM fibrosis (disease) and respective control conditions. We sort-purified Lin^neg^CD45^neg^tdTom^+^ cells from lineage-depleted BM and digested bone chip (BC) fractions (gating Figure S1A). An initial clustering after batch correction and integration identified 15 clusters, which displayed distinct levels of td-tomato expression (Figure S1B, S1C). We excluded cell clusters with low numbers of cells characterized as non-stromal cells (hematopoietic, neuronal and skeletal muscle cells) from further analysis resulting in a single cell experiment with 29,742 cells with mean genes per cell of 2,337. A high granularity re-clustering of the data recovered 30 clusters, which are associated with six major cell populations (Figure 1B): Cxcl12-abundant reticular cells (CAR cells; CAR1-7, IFN_CAR and MusCAR), fibroblasts (FB; FB1-8 and IFN_FB), osteo-lineage cells (OLC1-3 and osteoblasts; OB), chondrocytes (chondro and pre-chondrocytes), stromal cells (including smooth muscle cells), endothelial cells (Endo1; Endo2) including associated pericytes sub-populations. These are supported by the expression of *Cxcl12* (CAR), *Bglap* (OB), *Dcn* (FB) and *Pecam1* (Endo+pericytes) (Figure 1C, S1D).

Prompted by the specific spatial location of the Cre-reporters in imaging, we derived expression signatures from a recently published bulk-RNAseq dataset of laser-microdissected regions of homeostatic BM that define the endosteum-, sub-endosteum (marrow close but not adjacent to the endosteum), sinusoidal and arterial areas (16) to predict the spatial localization of the stromal cell clusters. The bulk-RNAseq-derived sinusoidal niche signature contained mainly hematopoietic genes, and was not retrievable in our deconvolution analysis as our data is enriched in stromal cells. Strikingly, and in line with the imaging, the Gli1;tdTom dataset was highly enriched in the arterial and endosteal signatures, the Pdgfrb;tdTom dataset confirmed the observation of wide-spread representation of Pdgfrb-lineage cells within the BM, and the Grem1;tdTom dataset was limited to the endosteal signature (Figure 1D). We thus recovered the ‘location stamp’ of the tdTom^+^ cells within the transcriptome of single cells.

### Fibroblasts expand in BM fibrosis while CAR cells are depleted after reprogramming and upregulation of collagen

To evaluate the cellular changes occurring during BM fibrosis, tamoxifen-mediated recombination of stromal Cre-reporters was induced in adult mice. In MPN, the Thrombopoietin receptor MPL is activated by the three driver mutations of MPNs (JAK2, CALR or MPL mutations) or can be activated by Thrombopoieitin. MPL is indispensable for MPN development regardless of the MPN driver mutation. The overexpression of thrombopoietin (TPO-OE), as a Mpl ligand, was used as a well-established model of myelofibrosis and osteosclerosis (17). ckit+ hematopoietic stem and progenitor cells (HSPCs) were lentivirally transduced with the TPO-OE plasmid or the respective empty vector (EV) control and transplanted into the lethally irradiated stromal Cre-reporter mice (Figure 2A). TPO-OE transplanted mice developed a robust phenotype of myeloproliferation reflected by high platelet- and white blood cell counts, and a drop in hemoglobin (Hgb) and BM cellularity due to reticulin fibrosis, which was accompanied by splenomegaly with extramedullary hematopoiesis (Figure S2A-D).

**Figure 2.**
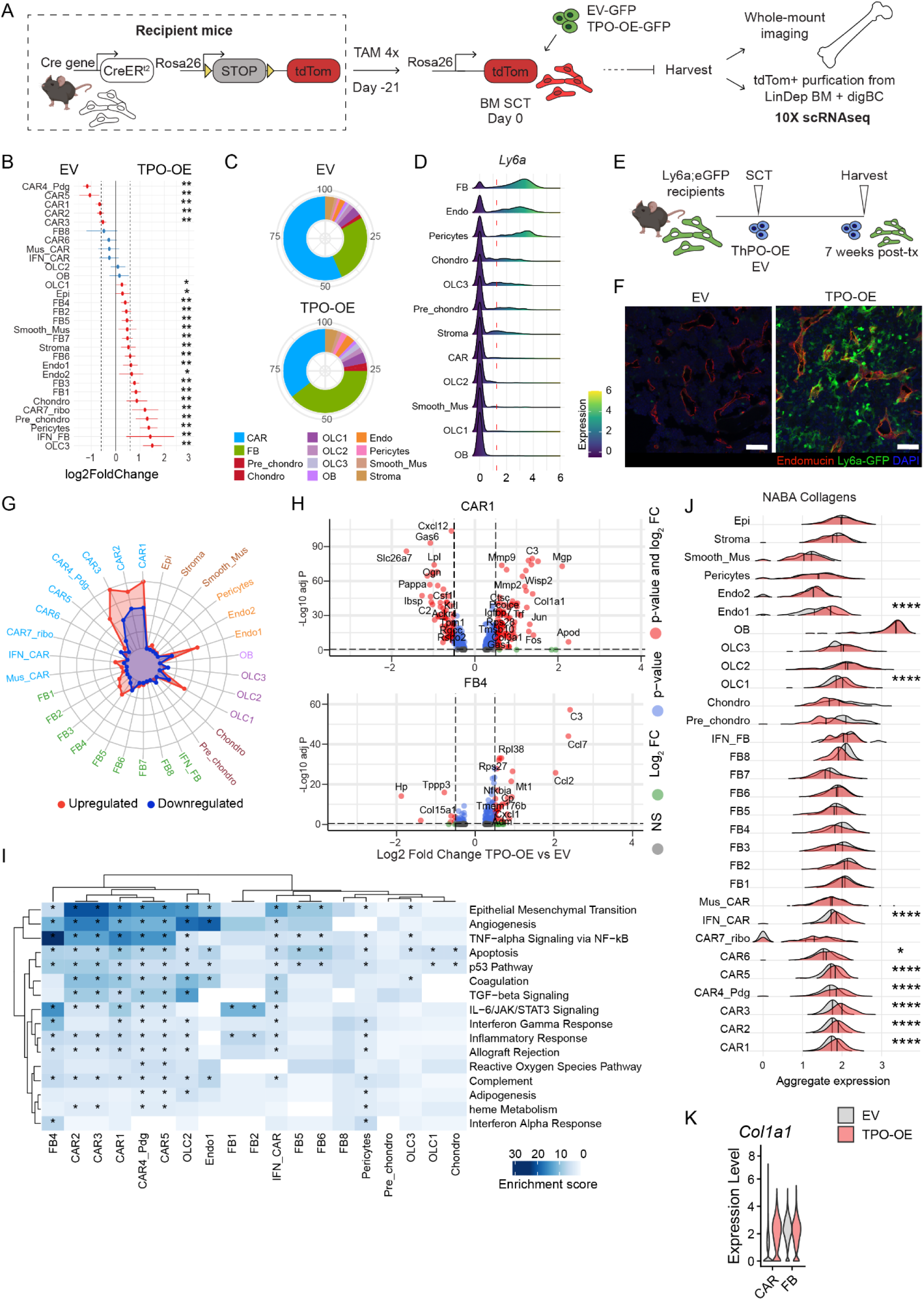
CAR cells acquire a pro-fibrotic phenotype but are reduced in frequency while fibroblasts expand in bone marrow fibrosis. (A) Experimental design: bigenic CreER;tdTomato mice were injected with tamoxifen (TAM), lethally irradiated at 21 days after the last tamoxifen dose, and intravenously received c-kit-enriched HSCs from WT littermates expressing either thrombopoietin cDNA (TPO-OE) or control (empty vector; EV) cDNA (both lentiviral SFFV-iGFP vector backbone). BM SCT = bone marrow stem cell transplant (B) Proportion test of cells per cluster (TPO-OE versus EV) obtained using scProportionTest. Relative differences in cell proportion per cluster, red colored dots show significant fold change (FDR<0.05 and absolute fold change > 0.58) with error bars showing confidence intervals for the magnitude difference (permutation test, n=1000). Blue dots: not significant; red dots: significant changes. (C) Contribution of individual clusters to all recovered cells per Cre-driver comparing the control (EV) to the fibrosis (TPO-OE) condition. (D) Ridge plot of *Ly6a* expression per cluster. Clusters are ordered regarding mean *Ly6a* expression values. (E) Ly6a;eGFP mice were lethally irradiated and intravenously received c-kit-enriched HSCs from WT littermates expressing either thrombopoietin cDNA (TPO-OE; n = 5, three males) or control cDNA [empty vector, EV, n = 5; both lentiviral SFFV-iblue fluorescent protein (BFP) vector backbone] scale bar = 100µm. (F) Representative images of TPO-OE BM of Ly6a;GFP mouse, Endomucin in red, DAPI nuclear stain in blue, scale bar = 100µm. (G) Radarplot of number of differentially expressed genes per cluster (TPO vs EV) (H) Representative volcano plots (TPO-OE versus EV) of differentially expressed genes in CAR1 and FB4. (I) Heatmap of enrichment scores using EnrichR of the up-regulated differentially expressed genes(TPO vs EV), genes with p-value < 0.05 were considered. Significance of the enrichment test was obtained using Fisher’s exact test implemented by EnrichR (J) Aggregate expression of NABA Collagens for all clusters (gene sets from (21). One-sided Wilcox test compared the expression of EV and TPO. (K) Collagen1a1 (*Col1a1*) expression in CAR cells and fibroblasts (FB) in TPO-OE versus EV control. Statistical significance is indicated by: *p<0.05, **p<0.01, ***p<0.001, ****p<0.0001.

ScRNAseq of tdTom+ cells from all Cre-reporters demonstrated that CAR cells were the most abundant tdTom+ stromal cell type in the control condition (EV) in line with the literature and their central role in hematopoiesis-support (Figure 2B, C). In fibrosis (TPO-OE), OLC-, FB- and chondrocyte-like cells were enriched within the sorted tdTom^+^ fractions, while all CAR cell populations were less represented. This is a fibrosis-specific response, as after irradiation and transplant (EV condition), we saw an increase of CAR and endothelial cells compared to the unperturbed niche (homeo) (Figure S2E). We were intrigued by the expansion of fibroblasts in BM fibrosis as fibroblast subsets were described as central players in solid organ fibrosis (3,18). In the BM, fibroblasts have been rather poorly defined in particular in response to hematological malignancy.

We thus explored our data for characterizing markers to discriminate fibroblasts from other stromal cells by applying sc2marker (19). *Ly6a* was ranked as the top marker for fibroblast identification (Figure 2D). *Ly6a* encodes for Sca1 glycoprotein, classically used to enrich hematopoietic stem cells from adult murine BM. Importantly, we did not observe an increase of *Ly6a* expression in the fibrotic marrow in non-fibroblast populations and endothelial cells, thus making it an attractive marker to study the fibroblast fate (Figure S2F). To validate that Ly6a^+^ fibroblasts expand in BM fibrosis, we used a Ly6a-GFP transgenic mouse strain (20) to label Ly6a^+^ cells upon fibrosis induction (Figure 2E). To avoid hematopoietic-derived Ly6aGFP signal, we performed a BM transplantation into Ly6a-GFP recipients with WT donor cells that were transduced with EV- or TPO-OE-BFP constructs (Figure 2E). The TPO-OE mice developed a typical myeloproliferative phenotype (Figure S2G-I). To account for endothelial cells being marked by *Ly6a* expression, we stained sections from the transplanted mice with the vascular marker endomucin (Emnc). In control conditions, Ly6a-GFP^+^Emnc^+^ cells were mostly found in perivascular localization, highlighting endothelial cells and pericytes (Figure 2F). In fibrotic BM, Ly6a-GFP^+^Emnc^-^ cells were massively expanded in the central marrow, partially in association to the vasculature but also in clusters of cells without direct vascular contact (Figure 2F).

We next asked how the expression profiles changed after induction of BM fibrosis. CAR cells had the highest number of differentially expressed genes (DEGs), followed by osteolineage cells (OLC1) and also a subcluster of endothelial cells (endo-1; Figure 2G). In contrast, fibroblasts as the most expanded cell population only showed minor transcriptional changes. CAR1 cells mainly upregulated collagen and ECM signature genes (*Col1a1*, *Col3a1*), matrix remodeling genes (matrix metalloproteinases *Mmp2*, *Mmp9*) and increased bone metabolism (*Apod*) (Fig 2H) (22). In line with previous findings (8), CAR1 cells decreased their hematopoiesis-supporting capacity (*Cxcl12*, *Lpl*, *Kitl*) (Figure 2H). The most significant up-regulation of pro-fibrotic TGF-beta signaling, as the master switch of fibrosis, occurred in CAR cell subsets, alongside an enrichment of epithelial-mesenchymal transition signatures (Figure 2I). BM-resident fibroblasts (here exemplary FB4 as population with highest DE genes) mainly gained an inflammatory signature, with upregulation of chemokines (*Ccl2*, *Ccl7*), components of the complement system (*C3*) (Figure 2H) and Nfkb-mediated induction of inflammation (Figure 2I). While CAR cells also gained a pro-inflammatory and increased apoptotic signature in the fibrosis setting, FBs in fibrosis were mainly characterized by a pro-inflammatory state known to play a role in the pro-fibrotic signaling and remodeling of the microenvironment.

Assessing the aggregate expression of collagen proteins (23) per cluster (Figure 2J), almost all CAR cell subsets showed an upregulation of collagen in exposure to TPO-OE hematopoietic cells. Interestingly, the transcriptionally most active subsets of osteolineage cells (OLC1) and endothelial cells (endo-1) also upregulated collagen expression. It is important to mention here that fibroblasts at baseline already have high expression of collagens and might, by expanding, also contribute to the fibrotic transformation of the BM (Figure 2K).

### Distinct stromal phenotype switch in metaphyseal and diaphyseal localization during fibrotic transformation

As the stromal Cre-reporters occupy distinct anatomical regions of the BM (compare Figure 1), we aimed to gain more spatial information on fibrosis formation by performing bone marrow 3D confocal microscopy (Figure 3; Supplementary figure 3).

**Figure 3:**
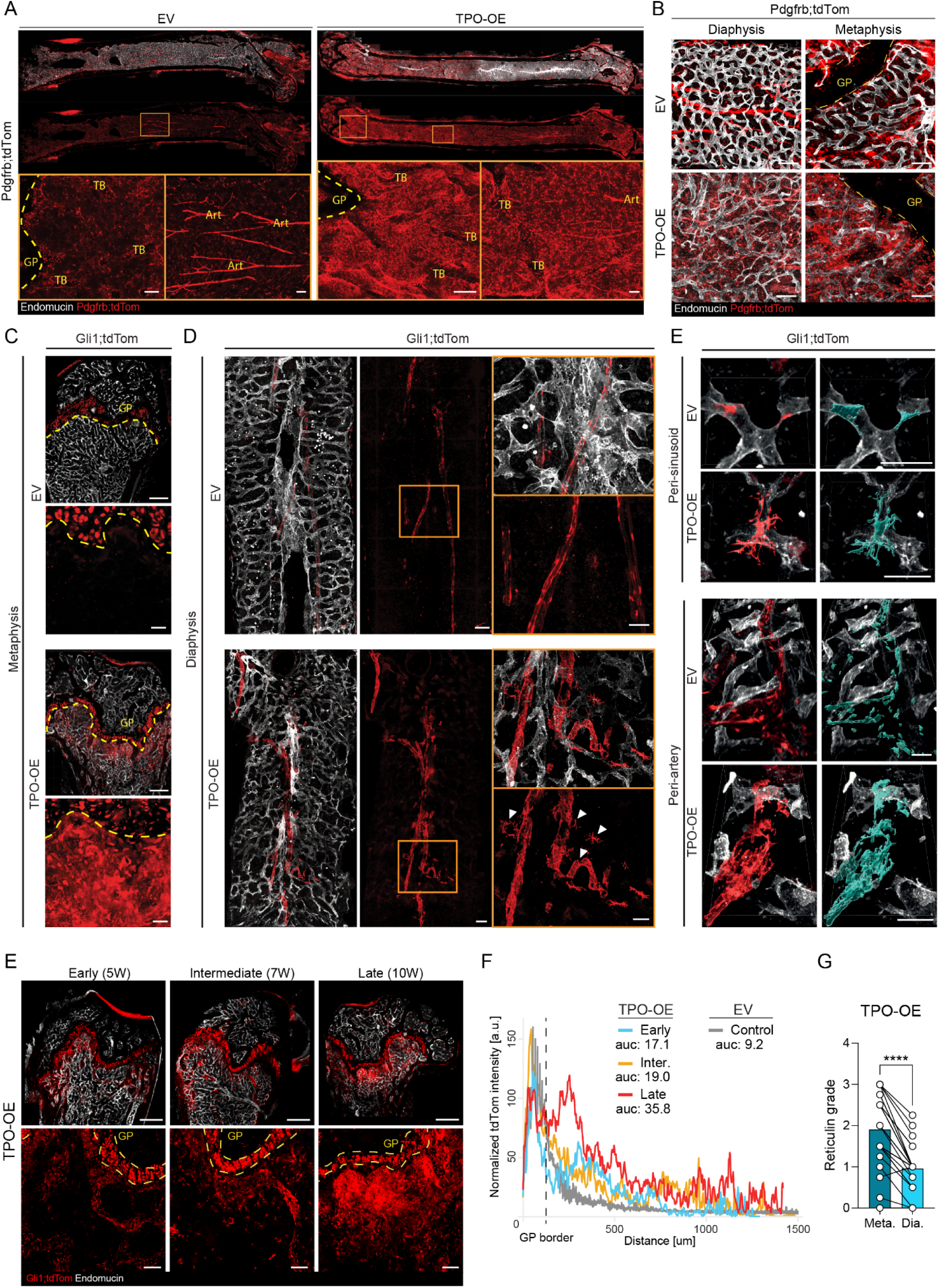
Distinct phenotype switch in TPO-OE induced bone marrow fibrosis in the diaphyseal and metaphyseal localization. (A) Representative whole-mount confocal images of Pdgfrb;tdTom femurs and zoom-in at indicated regions of the bone, EV= empty vector (control), TPO-OE = thrombopoietin overexpression (fibrosis). Art= arteries, TB = trabecular bone; dotted line demarcates the growth plate (GP). Scale bar = 100µm (B) Representative whole-mount confocal images of Pdgfrb;tdTom femurs at indicated regions of the bone, EV= empty vector (control), TPO-OE = thrombopoietin overexpression (fibrosis). Dotted line demarcates the growth plate (GP), Scale bar = 100µm. (C) Overview of whole bone of control (EV) and fibrotic Gli1;tdTom femurs, representative whole-mount confocal images. Dotted line demarcates the growth plate (GP). Scale bars = 300µm. Lower panel is zoom-in of metaphyseal region in EV and TPO-OE, respectively, scale bars = 20µm (D) Diaphyseal region of Gli1;tdTom bones, with Gli1-lineage cells wrapping around the central artery in control (EV) and fibrosis (TPO-OE) shown with and without endomucin co-staining. Boxes in the middle panel highlight the zoom-ins in the right panel. Note the detaching, large stromal cells leaving the arterial walls in the fibrotic setting highlighted by the white arrow heads. Scale bars middle panel = 100µm, scale bars zoomed in panels= 50µm (E) Morphological switch of Gli1;tdTom+ cells in peri-sinusoidal and peri-arterial location in fibrosis (TPO-OE) compared to control (EV). Representative whole-mount confocal images of single cells shown. Mask of Gli1;tdTom channel shown in blue. Scale bar = 50µm (F) Whole-mount confocal visualization of time-course Gli1;tdTom+ cell expansion after transplant with TPO-OE cKit-enriched cells. Early = 5W post-Tx, intermediate = 7W post-Tx, late = 10W post-Tx. Scale bar, top panel: 500µm, bottom panel: 50µm. Dotted line demarcates the growth plate (GP). (G) Quantification of Gli1;tdTom+ cell expansion/movement away from growth plate (GP) border from panel (F). Inter. = intermediate, auc = area under curve from the GP border, measured per line profile shown (H) Reticulin grading of fibrosis according to WHO criteria in the metaphysis (meta) compared to the diaphysis (Dia). Statistical significance is indicated by: *p<0.05, **p<0.01, ***p<0.001, ****p<0.0001.

tdTom^+^ cells from all Cre-reporters were activated by the presence of TPO-OE cells (Figure 3 and S3), but in distinct regions of the BM. While Pdgfrb;tdTom^+^ cells expanded in the diaphyseal region in BM fibrosis, the largest expansion was seen in the metaphysis, surrounding trabecular bone (Figure 3A, B). Within the metaphysis of fibrotic marrows, we observed signs of osteosclerosis, highlighted by Pdgfrb;tdTom+ cells expressing podoplanin, and their osteo-lineage-like morphology (Figure S3A). The Grem1;tdTom reporter was specifically detected in the metaphysis marking chondrocyte columns in the growth plate but Grem1;tdTom1^+^ cells showed only subtle changes after exposure to TPO-OE hematopoietic cells. Sparse, reticular-shaped Grem1;tdTom^+^ cells started to emerge from the growth plate (Figure S3B, arrowheads).

Gli1;tdTom^+^ stromal cells showed two distinct activation patterns upon fibrotic activation: increased frequency and expansion in the endosteal region of the metaphysis, and detachment from vasculature in the diaphysis (Figure 3C). In the diaphysis, Gli1;tdTom^+^ stromal cells increased in size, and gained an activated, myofibroblast-like morphology with long cellular extensions compared to cells in the control setting (Figure 3D, E). In these areas, endothelial cells also appeared activated, most likely due to the detachment of tdTom^+^ stromal cells from the vascular wall, leaving endothelial cells unprotected and exposed to the cytokine- and growth factor-rich environment caused by the fibrosis-inducing hematopoietic cells. The vascular bed in fibrotic marrow generally became more disorganized (compared Figure 3A-E; S3B). This is in line with transcriptional changes found in the Endo-1 cluster detected in the scRNA-seq (Figure S3C). This cluster showed upregulation and expression of apelin only in fibrosis, which is a marker for H-vessels, and an increase in endothelial-to-mesenchymal transition (endoMT; Figure S3D); (24).

Thus, the most distinct hotspot of tdTom^+^ stromal cell expansion in BM fibrosis in all Cre-reporters was within the metaphyseal region (Figure 3A-C, S3A-B; S3E). Spatiotemporal resolution combining a time course experiment with confocal imaging demonstrated the activation of Gli1;tdTom^+^ stromal cells in the metaphysis in a stepwise manner (Figure 3F). In the early phase (low grade fibrosis), spindle-shaped cells emerged from the trabecular bone of the metaphysis and increased over time into the central marrow. In progressed fibrosis (late), dense nests of Gli1;tdTom^+^ cells were found broadly beneath the growth plate with activated morphology. To assess the activation of Gli1;tdTom^+^ cells from metaphysis to diaphysis, we quantified the progression of tdTom-signal from the growth plate (GP) to central marrow (Figure 3G). Here, we found a stepwise increase in the tdTom signal in fibrotic marrow compared to EV controls, with increases of tdTom signal present even in early-stage disease. We included an additional model of BM fibrosis by transplanting cKit+ cells transduced with the MPLW151L (MPL) as a common MPN and MF-driver mutation (6) into Gli1;tdTom recipient mice. It is important to note that the retroviral MPL murine model is characterized by strong myeloproliferation but less severe fibrosis compared to the TPO-OE model (compare fibrosis grades Figure 3H and S3F) (6). We observed a notable increase in Gli1;tdTom+ mobilization in the more fibrotic TPO-OE marrows, unlike in MPL-transplanted marrows (Figure S3G), indicating that advanced fibrosis is associated with enhanced Gli1;tdTom+ expansion into the BM. We overlaid a reticulin staining on the imaged tdTomato signal after demounting of slides in the Pdgfrb-reporter (Figure S3E). Interestingly, the tdTom+ hotspots completely aligned with the hotspots of reticulin deposition in the metaphyseal region, while reticulin deposition in the diaphyseal region of the same bone was rather minor (Figure S3E). In line with this, we quantified reticulin fibrosis in the metaphysis and diaphysis of the same bone, and found significantly higher fibrosis in the metaphyseal region in both the TPO-OE (Figure 3H) and MPL models (Figure 3H and S3F). This data suggests that an injury-specific stromal population resides within the metaphysis which is essential for the fibrotic transformation.

### OLC3 represent stromal progenitor cells and reside in the metaphysis of the bone marrow

We hypothesized that the metaphyseal (trabecular bone) and diaphyseal (compact bone) regions contain stromal cells with distinct functions and differentiation potentials, in line with reports on skeletal repair (25). Specifically, we asked if the peritrabecular region is a reservoir for stromal and fibrosis-driving cells, which react to injury/inflammatory stimuli.

We thus sought to 1) establish a stromal cell hierarchy and 2) assess the lineage trajectories and cell fates of these populations. We applied PHATE visualization reduction of the data, focusing on disease-relevant cell types with significant numerical or transcriptional changes: CAR cells, FBs, OLCs, chondrocytes, and terminally differentiated osteoblasts (Figure 4A, compare to Figure 2). In this projection, OLC3 cells were positioned as a central hub connecting CAR cells, pre-chondro- and chondrocytes and fibroblasts (Figure 4A). In a different PHATE projection, OLC3 were positioned at the apex of the differentiation hierarchy, demonstrating connections to the CAR4_Pdg cluster, the pre-chondro and FB clusters, as well as the mature OBs and chondrocytes (Figure S4A).

**Figure 4.**
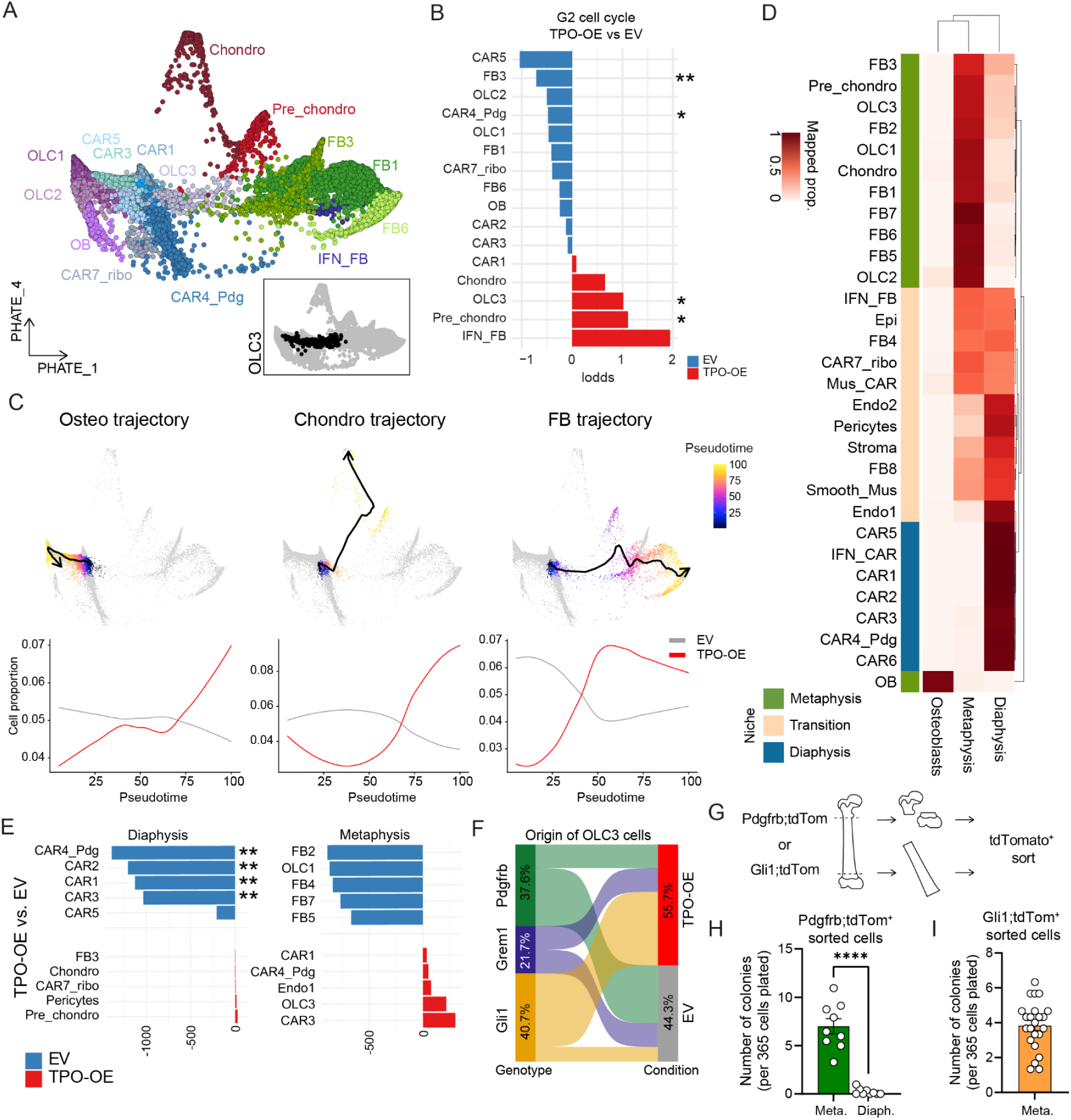
OLC as progenitor cells of the bone marrow are enriched in the metaphysis in bone marrow fibrosis. (A) PHATE dimension reduction of CAR, OLC and FB cells indicate potential cell differentiation trajectories. Inset in lower right corner showing PHATE visualization with OLC3 cluster highlighted in black. (B) Cell cycle G2 proportion analysis in stromal cell clusters, comparing TPO-OE vs. EV control. Statistical comparison was done using Fisher Exact Test. (C) Trajectory analysis of CAR cells/OLCs towards Osteo, Chondro and FB cells (top). Proportion of cells in EV or TPO conditions along pseudotime (bottom) per experimental condition (D) Proportion of cells assigned to ‘diaphysis’, ‘metaphysis’ and ‘osteoblast’ reference cell dataset (source Srivaja et. al. dataset). (E) Difference in the absolute proportion of cell clusters associated with diaphysis and metaphysis in TPO vs EV conditions. Asterisks represent cells with differences larger than two standard deviations around the mean. (F) Sankey plot indicating the origin of OLC3 cells, per genotype and condition. (G) Experimental setup on the isolation of cells from the metaphyseal (peritrabecular) compared to the diaphyseal region of homeostatic Cre-reporter lineage traced bones. (H) Quantification of fibroblastic colony-forming units (CFU-F) of sort-purified Pdgfrb;tdTom+ cells (n = 3 biological replicates, 3 technical replicates, data are presented as mean +/- SEM, two-tailed unpaired t-test). Meta. = metaphyseal region, Diaph. = diaphyseal region. (I) Quantification of fibroblastic colony-forming units (CFU-F) of sort-purified Gli1;tdTomato+ cells (n = 4 biological replicates, 5-6 technical replicates, data are presented as mean +/- SEM). Meta.= metaphyseal region, Diaph. = diaphyseal region Statistical significance is indicated by: *p<0.05, **p<0.01, ***p<0.001, ****p<0.0001.

To achieve an unbiased view of the progenitor and expansion potential of the stromal clusters, we explored both the coefficient of variance score of cells (as surrogate marker for stemness or specialization), as well as their cell cycle status. In brief, cells with a higher coefficient of variance have highly specialized expression of genes, termed ‘SPEC’, supporting a differentiated state. In contrast, cells with a lower coefficient of variance, termed ‘MONO’ or monotonous, do not display any specific expression pattern as typically seen in progenitor-like cells. The application of these scores, suggested that CAR4_Pdg is a stem and progenitor cell population (low SPEC and high MONO) and the osteoblast- and chondrocyte clusters scored highly as differentiated cells (high SPEC and low MONO);(Figure S4B). Next, we concentrated on the cell cycle state of cells. We observed a significant increase in the G2 cell cycle stage in OLC3 and pre-chondro populations in TPO-OE vs. EV, and a reduction of these populations in G1 cell cycle stages (Figures 4B and S4C). OLC3 cells populations also displayed an intermediate stemness (MONO) and a low specialization (SPEC) score. This indicates a progenitor role of the OLC3 cluster, while its proliferation in fibrosis suggests the activation in response to tissue injury.

We used this information to estimate trajectories and pseudotime using ArchR and PHATE (Figure 4C). In PHATE visualization, three distinct trajectories appeared: 1) an “osteo-primed trajectory” through CAR1/3/5, OLC1/2/3 towards mature OBs, 2) an “chondro-primed” trajectory via OLC3 towards chondrocytes and 3) a “FB primed” trajectory through OLC3 towards FBs. We next ran pseudotime analysis along the trajectories. Cells showed an increase of specialization (decrease of stemness) scores in pseudo-time (Figure S4D). The osteo-, chondro- and FB- trajectories were increased in BM fibrosis (TPO-OE). The increased differentiation into fibroblasts indicates that their abundance in the BM in fibrosis can be explained by activation of progenitor cells.

To explore the spatial relevance of fibrosis deposition in the BM, we mapped scRNAseq clusters to their spatial localization in the BM, using a published dataset defining regional specifications of metaphyseal and diaphyseal BM stromal cells.(25) Label transfer analysis showed that most CAR clusters, including CAR4-Pdg, were associated with a diaphyseal signature, while some had a mixed location, likely representing a transition zone (Figure 4D and 4E). Metaphyseal signatures were enriched in chondrocytes, FBs, and OLCs -the cell types predominantly found in TPO-OE BM (Figure 2B). Comparing TPO-OE to EV metaphyseal and diaphyseal signatures, we observed a loss of diaphyseal-location signature in CAR clusters, suggesting perivascular detachment and fibrotic reprogramming (Figure 3). Osteo-primed CAR3 and OLC3 showed an increase in metaphyseal signatures, indicating peritrabecular expansion, while the metaphyseal signature of FBs and OLC1 decreased in TPO-OE. Ly6a+-fibroblasts, typically absent in the diaphysis at steady state, were abundant in fibrosis (Figure 2F).

We thus wondered how the different Cre-drivers contribute to OLC in control (EV) and fibrosis (TPO-OE). While under control conditions, Pdgfrb;tdTom+ cells mostly contribute to OLCs, these subsets are mostly derived from Gli1;tdTom+ cells in fibrosis confirming the fibrosis-specific activation of Gli1+ cells (Figure 4F).

To functionally validate the stem and progenitor potential of cells in the metaphyseal region, we sort-purified tdTom+ cells from the metaphyseal and diaphyseal regions of long-bones of unperturbed, homeostatic mice (Figure 4G). By applying the Pdgfrb;tdTom reporter for broad stromal lineage marking, we saw comparable tdTom+ fluorescent signal strengths between metaphyseal and diaphyseal tdTom^+^ cells, with similar frequencies of tdTom^+^ cells between the macroniches (Figure S4E). Remarkably, only tdTom^+^ cells isolated from the metaphysis regions of long bones harbored colony forming potential, and diaphyseal tdTom+ cells from the same bones did not give rise to fibroblast colonies *in vitro* (Figure 4H). Gli1;tdTom cells were mainly located in the metaphyseal region in homeostatic mice (Figure S4F). Sorted Gli1;tdTom+ cells from the metaphyseal region contained a high number of CFU-Fs, supporting their progenitor-cell state (Figure 4I).

To validate that Pdgfrb;tdTom^+^ and Gli1;tdTom^+^ cells can mark long-lived mesenchymal progenitor cells, we induced recombination in adult mice and harvested bones more than one year after the last tamoxifen injection (Figure S4G). Pdgfrb;tdTom^+^ cells were distributed across all compartments of the bone (metaphysis, diaphysis), whereas Gli1;tdTom^+^ cells were mainly positioned in the metaphysis and less present in the diaphysis. Comparing the Pdgfrb;tdTom and Gli1;tdTom reporters in homeostatic conditions revealed that the Pdgfrb-reporter is enriched for CAR4_Pdg cells (stromal stem cell). In contrast, Gli1^+^ cells are mainly enriched for the OLC3 population (Figure S4H). We thus propose that PdgfrbCre;ERT2 labels a perivascular stromal stem cell (CAR4_Pdg) that replenishes the BM stromal population under physiological conditions, and Gli1Cre;ERT2 labels a peritrabecular mesenchymal progenitor cell population (OLC3) that is activated in an injury-specific manner from their peritrabecular localization.

### CAR and OLCs are skewed in their differentiation in BM fibrosis

We hypothesized that the fate of the peritrabecular OLC3 and perivascular CAR4_Pdg stem and progenitor cells is skewed in the presence of fibrosis-inducing hematopoietic cells and an inflammatory environment. We indeed observed distinct histomorphological changes in fibrotic BM compared to non-fibrotic controls. After irradiation, control BM (EV) typically depict adipocytes in the metaphyseal region (Figure 5A). In contrast, fibrotic marrows were almost devoid of adipocytes (Figure 5A). This change was quantifiable using MarrowQuant (26). Non-hematopoietic and non-vascular cellular components were expanded in fibrotic (TPO-OE) marrows while the adipocyte area and number of adipocytes decreased (Figure 5B, C).

**Figure 5:**
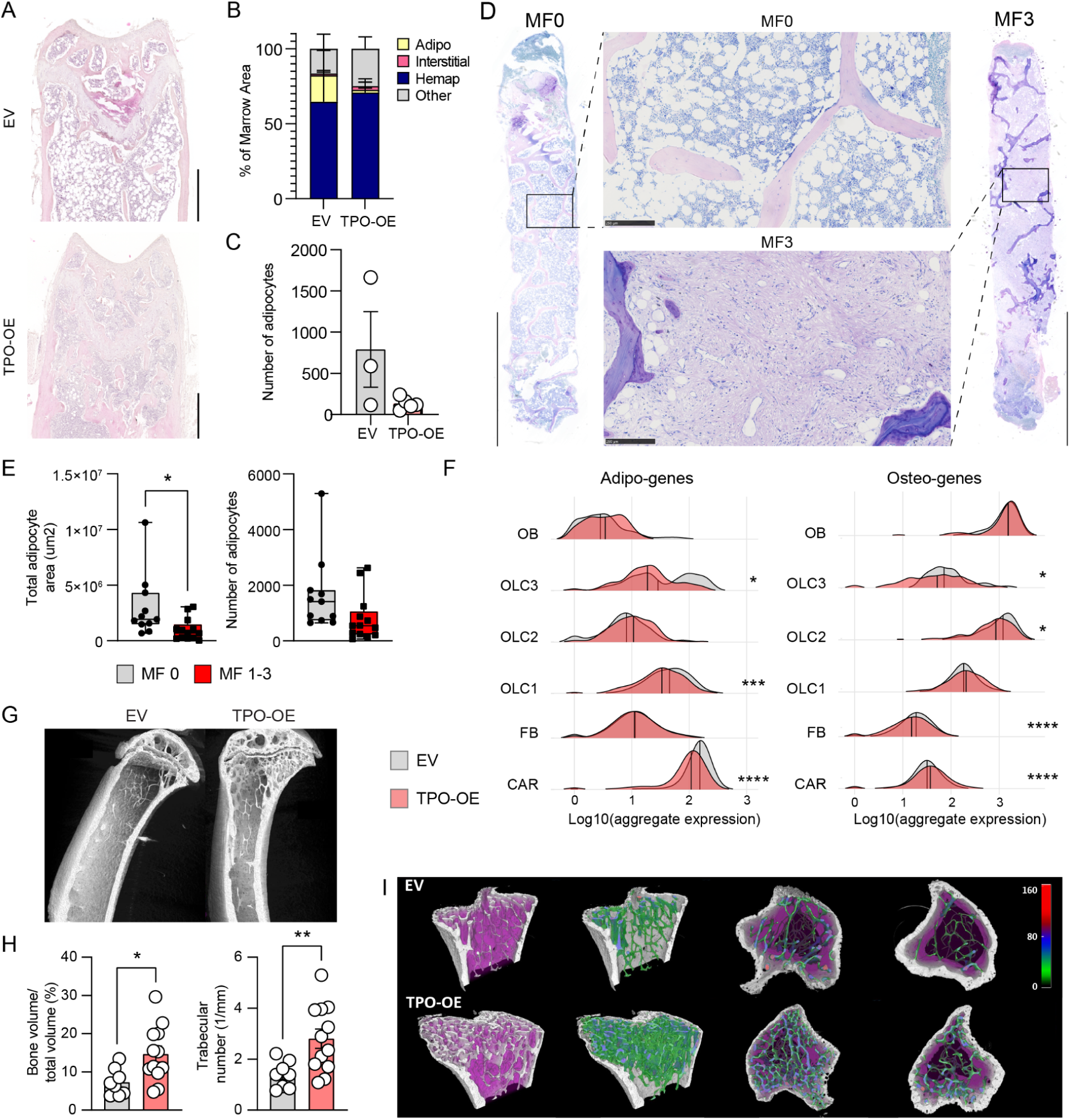
Stromal stem and progenitor cells are skewed in their differentiation in bone marrow fibrosis. (A) Representative images of HE staining of wild-type recipients transplanted with EV or TPO-OE BM 7 weeks after transplantation. Scale bar = 500µm. (B) Analysis of bone marrow composition using Marrowquant in H&E stained EV and TPO-OE femurs. EV n=3, TPO-OE n=5. (C) Number of adipocytes in H&E stained EV and TPO-OE femurs. EV n=3, TPO-OE n=5, t-test for significance. (D) Representative overviews of H&E stained biopsies from MPN patients with myelofibrosis grade 0 (MF0) and MF3. Scale bars = 1mm (overview image), 250µm for zoom-in (E) Quantification of total adipocyte area and number of adipocytes in cohort of MPN patients with varying grades of fibrosis (n=11 MF0; n=13 MF1-3), unpaired t-test significance test, * p<0.05 (F) Ridge plot showing aggregate expression of adipo- and osteo-genes per cluster, (TPO vs EV), two-sided Wilcox test. (G) Representative capture of µCT of tibia in mice transplanted with EV or TPO-OE overexpressing bone marrow 7 weeks post transplantation; scale bar = 1mm. (H) Quantification of trabecular bone from µCT bones specifying bone volume over total volume and trabecular number. (I) Representative 3D reconstructions of tibia in mice transplanted with EV or TPO-OE bone marrow with trabecular thickness quantification shown. Color code represents the trabecular thickness in µm. Statistical significance is indicated by: *p<0.05, **p<0.01, ***p<0.001, ****p<0.0001.

The pattern of decreased adipocytes in fibrosis was confirmed in other murine models of myeloproliferation associated with myelofibrosis (MF): JAK2V617F- and MPLW515L-mutated disease (Figure S5A). Importantly, the decrease in adipogenesis was even confirmed in BM biopsies of myeloproliferative neoplasm (MPN) patients and specific to MPN with fibrosis, not MPN per se (Figure 5D, E), indicative of skewed differentiation of mesenchymal progenitor cells in fibrosis.

Given this apparent change in stromal differentiation in the context of BM fibrosis, we explored the expression of adipocyte (adipo), osteoblast (osteo) (25) and chondrocyte (chondro) (27) specific-genes within our scRNAseq dataset (Figure 5F and S5B, gene sets in table S2). The average expression of these genes per cluster grouped the homeostatic dataset into more adipo-primed (CAR cells) and osteo-primed cell clusters (OLCs and OBs); (Figure S5C). CAR3 were, however, more osteo-primed as indicated by relatively higher expression of *Spp1*, *Alpl*, *Sp7* and *Wif1* compared to other CAR populations. OLC1 is a predominantly osteo-primed population, but still expresses *Foxc1*, *Gas6* and *Apoe*. In fibrosis, a distinct skewing in the differentiation was observed: CAR cells, OLC1 and OLC3 significantly decreased their expression of adipo-genes, while specifically CAR cells increased the expression of osteo-genes (Figure 5F). The skewed differentiation was also obvious in chondro-linked genes in the fibrotic setting, specifically in the CAR populations and OLC3, potentially hinting at calcification occurring in these cells upon fibrotic transformation (Figure S5B). The increase of osteo-associated genes in fibrotic marrows was also confirmed by µCT analysis (Figure 5G-I). Bones in the fibrotic conditions had more bone, more trabeculae and more trabecular connectivity, compared to the control (EV) condition (Figure 5H).

### NCAM is a marker for fibrosis-driving cells in bone marrow fibrosis

We next applied sc2marker to identify specific markers to discriminate OLCs from fibroblasts (FBs) and CAR cells. The top markers identified were Fibulin (*Fbln1*) and *Ncam1* (CD56); (Figure 6A; Figure S6A). However, *Fbln1* represented a less specific marker as it was expressed to some extent in FBs and CAR cells as compared to *Ncam1* which was rather limited in its expression to OLCs and osteoblasts (Figure S6A). In line with this, confocal imaging revealed a more unspecific pattern for *Fbln1* expression (Figure S6B). *Fbln1* was predominantly expressed in spindle-shaped cells with perivascular localization and diffusely beneath the growth plate in early stages of fibrosis, with increased expression surrounding Gli1;tdTom+ cells in more progressed fibrosis. This might be explained by the fact that Fbln1 is known to integrate into fibronectin-containing matrix fibers, facilitating cell adhesion and migration along extracellular matrix (ECM) protein fibers, thus indicating lower specificity to mark OLCs (28).

**Figure 6.**
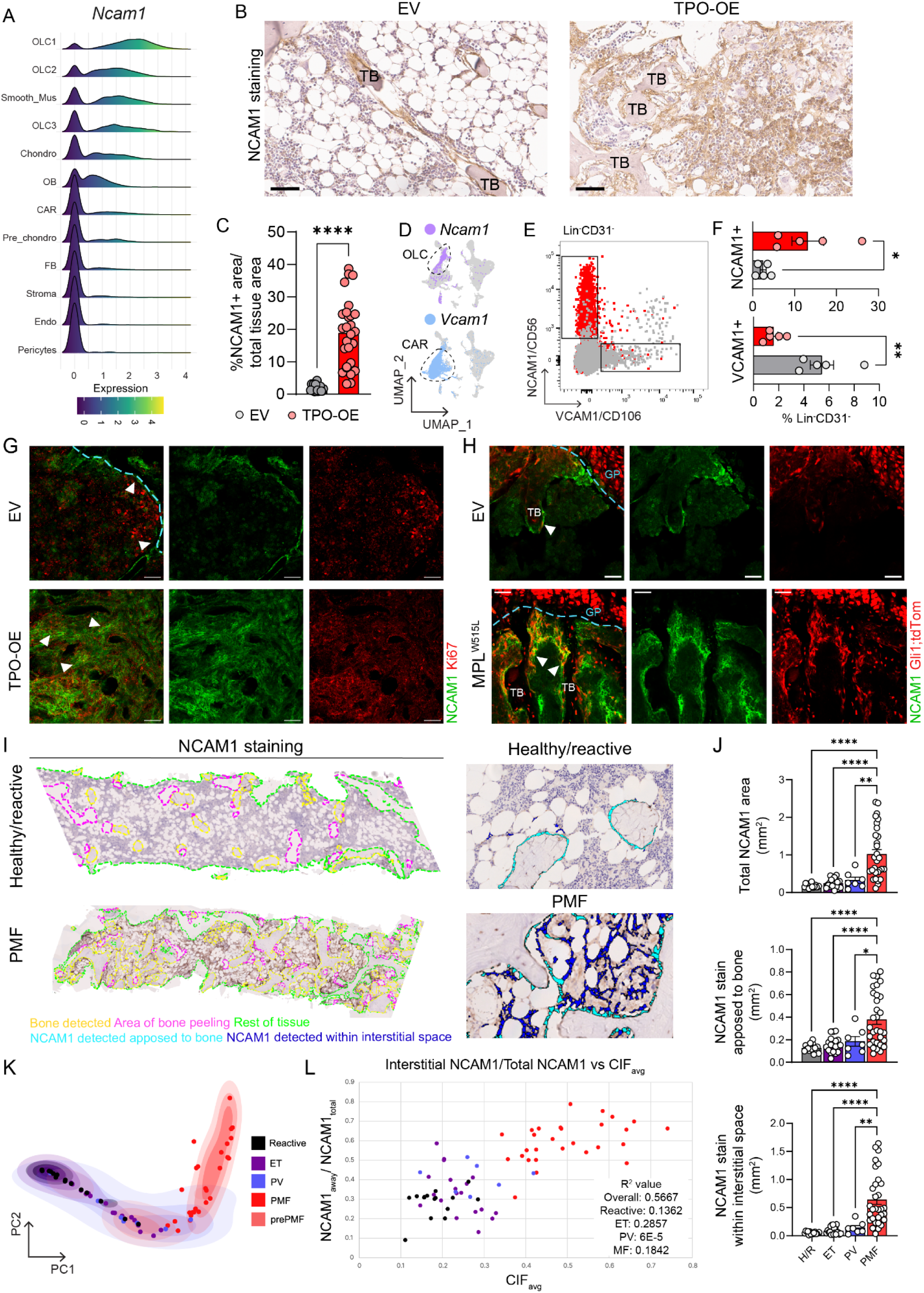
NCAM is a marker for fibrosis-driving cells in bone marrow fibrosis. (A) Ridge plot of *Ncam1* expression for clusters. Clusters are ordered by mean *Ncam1* expression values. (B) Immunohistochemical staining of NCAM1 in murine biopsies from control (empty vector, EV) or fibrotic (TPO-OE) BM. Scale bar: 50µm (40x).TB = trabecular bone (C) Quantification of NCAM1/CD56 staining in murine sections, one point represents the average stained area of three areas in murine bone, EV n=5, TPO-OE n=9, unpaired t-test (D) UMAP showing cells expressing *Ncam1* > 2 and *Vcam1* > 2, dotted lines related to global OLC and CAR clusters respectively. (E) Representative FACS plots of Lineage negative, CD31 negative BM and digested BC from transplanted mice at harvest, NCAM1/CD56 and VCAM1/CD106 gates shown. (F) Quantification of NCAM1+/CD56 and VCAM1+/CD106 populations from cohort in panel 6E. Unpaired t-test. (G) Representative confocal imaging of murine humeri, NCAM1/CD56 in yellow, Ki67 proliferation marker in red, scale bar = 50µm. (H) Representative confocal imaging of murine humeri in Gli1;tdTom reporter, NCAM1/CD56 in yellow, Gli1;tdTom in red, scale bar = 50µm (I) Overview of healthy/reactive and PMF patient bone marrow biopsy with annotated areas as indicated (left). The images on the right show in magnification the detected and quantified areas. (J) Quantification of NCAM in human biopsies as seen in 6I. Healthy/reactive (H/R) n=15, essential thrombocythemia (ET) n=19, polycythemia vera (PV) n=7 and primary myelofibrosis (PMF) n=31 samples. One-way ANOVA with multiple comparisons. (K) Principal component analysis (PCA) plot of the whole cohort using average image tile CIF score, tile distribution, and heterogeneity of tile distribution as variables, as previously described (29). Association between CIF (x-axis) and NCAM1/CD56 (y-axis) cell expansion (defined by NCAM1 staining in interstitial space/total NCAM1 staining). Statistical significance is indicated by: *p<0.05, **p<0.01, ***p<0.001, ****p<0.0001.

To validate *Ncam1* as a marker for peritrabecular progenitor cells on the protein level, we stained murine BM for NCAM1 (Figure 6B). In the control (EV) condition, NCAM1+ cells had a distinct localization just below the growth plate, seaming the osteogenic front towards the central marrow, in addition to surrounding trabeculae as spindle-shaped cells. In fibrosis (TPO-OE), NCAM1^+^ cells significantly expanded from their peritrabecular localization (Figure 6C). In dense centers of NCAM1^+^ cells, osteosclerosis was observed and adipocytes were decreased, in line with the observed skewed differentiation of OLCs as peritrabecular mesenchymal progenitor cells.

Sc2marker further identified *Vcam1* (CD106) as a good discriminatory marker for CAR cells (Figure S6C). The combined use of NCAM1 (CD56) and VCAM1 (CD106) allows for clear discrimination between OLC-enriched and CAR-enriched populations, as these markers display distinct expression patterns as projected on the UMAP (Figure 6D). Flow cytometry confirmed that NCAM1 and VCAM1 can be used to differentiate between distinct stromal cell populations, and that the abundance of NCAM1+ OLCs is increased in fibrosis, whereas there is a loss of VCAM1+ CARs (Figure 6E, F). Flow cytometry further showed that NCAM1+^+^ cells are specifically enriched in the metaphysis while VCAM1^+^ cells are ubiquitously distributed (Figure S6D, E). To characterize these distinct populations better, we sort-purified and immortalized them for characterization by FACS and qtRT-PCR *in vitro*. Both NCAM1^+^ and VCAM1^+^ cells express typical stromal markers, and express genes known to characterize stromal cells such as the hematopoiesis-support genes *Cxcl12* and *Kitl*, (Figure S6F-H). In co-culture experiments with ckit^+^ hematopoietic stem and progenitor cells (HSPCs), both NCAM1^+^ and VCAM1^+^ cells maintained a higher viability of ckit^+^ HSPCs and decreased apoptosis compared to cKit^+^ HSPC suspension culture without stromal support (Supplementary Figure 6F-J).

Having these discriminatory markers identified, we asked if the increased proliferation of NCAM1^+^ OLCs (compare Figure 4B and S4C) can be validated. Combined Ki67 and NCAM1 staining confirmed the increased expression of Ki67 in NCAM1+ cells lining the growth plate and located within nests of peritrabecular NCAM1+ cells in the metaphysis. In addition, we observed Gli1;tdTom^+^ NCAM1+ spindle-shaped proliferates detaching from the trabecular bone and expanding into the interstitial space in BM fibrosis (Figure 6H), highlighting the cellular origin of NCAM1+ OLCs in fibrosis.

To confirm the presence of peritrabecular OLCs in human MPN disease setting, we stained control and myelofibrosis (MF) patient samples for NCAM1 (Figure 6I). It is important to note that routinely taken diagnostic biopsies are from the pelvic region, which contains bone of trabecular quality. Using an automated digital pathology algorithm, we detected a significant increase in the overall area of NCAM1 staining specifically in primary myelofibrosis (PMF) patients (Figure 6J). In line with our murine data, NCAM1+ cells were located “apposed to bone” (i.e. peritrabecular location) in healthy individuals. Importantly, there were distinct NCAM1 distribution differences between the MPN entities. NCAM1^+^ cells were only found in high frequency in the central marrow (IT space) in the context of fibrosis in PMF but not in MPN in general. To measure the fibrotic content in the patients stainings, we computed the Continuous Indexing of Fibrosis (CIF) score and used the spatial distribution and heterogeneity of the CIF in individual patient slides (29) to obtain a principal components analysis (PCA) representation of our cohort (Figure 6K). The PCA discriminated patient samples of distinct entities and placed MF opposite to non-reactive (healthy) samples. Strikingly, when plotting the CIF score versus the “NCAM1 away from bone/NCAM1 total”, MF patients and pre-MF patients formed a distinct group. This validates the fact that the spatial distribution of NCAM1+ cells represents a solid marker for the different stages of fibrosis. It further highlights the potential of this readily available immunostain to identify patients in whom there is significant expansion / prominence of the NCAM1+ endosteal lining cells in the context of (osteo)myelofibrosis.

### Wnt signaling is upregulated in metaphyseal stromal progenitor cells and CAR cells

We next wondered if OLCs, as the peritrabecular progenitor cell reservoir for fibrosis-driving cells, can be therapeutically targeted. We thus investigated the molecular mechanisms that drive the activation of the stromal stem and progenitor cells in BM fibrosis with a focus on cell-cell interactions (CCI) using CrossTalkeR (30). As hematopoietic cells are the fibrosis-inducing cells and contribute to the inflammatory niche, we included hematopoietic cells in the analysis to not lose any disease-relevant signals (neutrophils 1, 2 and hemap progenitors). In line with our hypothesis that CAR cells in the diaphyseal localization are reprogrammed by the fibrosis-inducing hematopoietic cells, the most significant interactions occur between all CAR cell clusters and hematopoietic cells. Hematopoietic cells also showed increased interaction with fibroblasts, in line with their expansion throughout the marrow. IFN-CAR cells had significant interactions with OLCs, indicating that this subset is a main communicator in the fibrotic marrow (Figure 7A). We specifically looked into “chemokine” and “ECM regulator” interactions as drivers of BM fibrosis (Figure S7A, B). As expected, the most pronounced chemokine-driven interactions occurred in the diaphysis (perivascular niche) between hematopoietic cells and CAR cells, specifically IFN_CAR and CAR4_Pdg, but also to the metaphysis (trabecular niche) (OLC1/2). For the “ECM regulator”-dominated cross-talk, the main communication occurred between diaphysis and metaphysis, specifically between IFN_CAR and OLC2/3, and fibroblasts, stressing the essential role of these populations in the fibrotic transformation.

**Figure 7.**
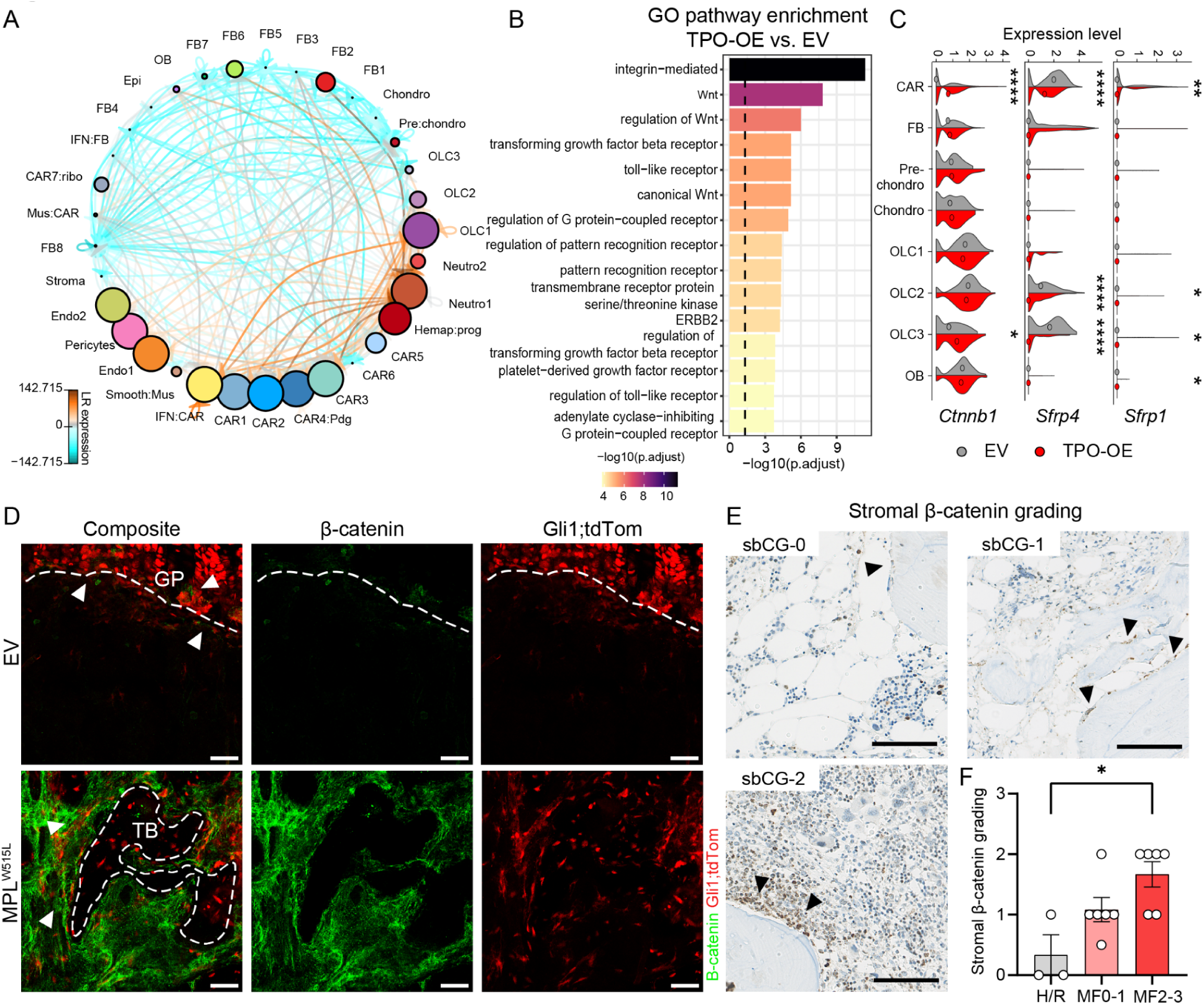
Wnt signaling is activated in OLC3 in bone marrow fibrosis. (A) CCI plot signaling showing ligand-receptor (LR) interactions of the different niches (metaphysis, diaphysis, transition vs. hematopoietic=hemap) and cell clusters (TPO-OE vs. EV). The size of a cell indicates its importance in the CCI network. The thickness of the edges indicates the number of LR interactions, while the color indicates the relative change in expression (orange is higher in TPO and blue is higher in EV). (B) GO enrichment of the enriched LR pairs with expression in TPO-OE versus EV in OLC3 cluster using ClusterProfiler (C) Violin plot depicting the Wnt modulator *Ctnnb1* and inhibitors *Sfrp1* and *Sfrp4 ,* median depicted with circle, wilcox test for significance, only significant results shown (D) Beta-catenin staining of murine humeri, EV= empty vector, control, MPLW515L = fibrotic marrow. Composite image shown, and the right panels show individual channels of beta-Catenin and Gli1;tdTom respectively. White arrowheads highlight beta-catenin and Gli1;tdTom co-expression. Scale bar = 50µm (E) Representative immunohistochemical staining of b-Catenin, each panel showing stromal beta catenin grade (sbCG) 0-2. Scale bar: 100µm (F) Grading of stromal beta catenin grading (sbCG) of human bone marrow biopsies, One-way ANOVA with Kruskal-Wallis. H/R = healthy/reactive, MF = myelofibrosis/reticulin grade 0-3. Statistical significance is indicated by: *p<0.05, **p<0.01, ***p<0.001, ****p<0.0001.

Due to the importance of OLC3 as a metaphyseal progenitor cell, we focused on OLC3 related receptor-ligand interactions and performed GO pathway analysis (Figure 7B). We identified central disease-relevant pathways in BM fibrosis such as integrin-mediated-, toll-like receptor-, platelet-derived-, and TGFb-receptor signaling, and particularly a distinct Wnt pathway signature. Filtering the CCI-plot for Wnt-pathway receptor-ligand pairs, Wnt-specific upregulation in the communication was noted in OLCs, CAR cells and fibroblasts (Figure S7C). Specifically in fibrosis, OLC3 and broadly CAR cells showed a significant increase in the expression of the intracellular signal transducer of the Wnt pathway, β-Catenin (*Ctnnb1*), as well as a decrease in expression of Wnt inhibitors *Sfrp1* and *Sfrp4* (Figure 7C).

Before exploring the relevance of targeting Wnt signaling in BM fibrosis, we aimed to validate the increased β-Catenin signature in CAR cells and OLC3 in Gli1;tdTom^+^ cells in BM fibrosis induced by the MPLW515L mutation. While in control BM, β-Catenin expression was mostly limited to either cells lining the growth plate (GP; Figure 7D), or to chondrocytes of the GP (Figure S7B), the expression increased in expanding Gli1;tdTom+ cells in the metaphysis in BM fibrosis (Figure 7D, S7E). We next asked if Wnt/β-Catenin is differentially expressed in the bone marrow stroma in patient samples. In control bone marrow biopsies, β-Catenin expression was limited to hematopoietic cells while spindle-shaped bone lining cells did not stain positively (Figure 7E). We termed this stromal β-Catenin grade 0 (s-bCG0; Figure S7F). In low grade fibrosis, almost all bone-lining cells expressed β-catenin (s-bCG1). In progressing fibrosis, all bone lining cells expressed β-Catenin and started to extend from the bone (s-bCG2). The grading of the s-bCG confirmed a significant increase of the s-bCG with progressed fibrosis, although interestingly the overall BM expression of β-Catenin, also taking hematopoietic cells into account, rather showed a decrease with MPN and fibrosis progression (Figure S7F).

### Wnt signaling activation in stromal compartment is a druggable target and its inhibition ameliorates bone marrow fibrosis

Given that the Wnt pathway is specifically active in peritrabecular mesenchymal progenitor cells that are activated in fibrosis, we tested the effect of Wnt inhibition on fibrosis and osteosclerosis. Canonical Wnt signaling orchestrates skeletal regeneration upon injury (31,32), so we hypothesized that its inhibition would keep OLCs uncommitted, halting their activation cycle. To this end, we repurposed Wnt inhibitors from cancer research, focusing on stabilizing the β-catenin destruction complex. We identified the FDA-approved drug Pyrvinium-Tosylate (PT), which activates casein kinase 1α (CK1α), thereby promoting β-catenin degradation through phosphorylation, ubiquitination, and proteasomal pathways.

To provide proof-of-concept that the inhibition of the Wnt pathway with PT has therapeutic and translational relevance in the fibrotic transformation, we started twice-weekly treatment of mice under fibrotic (TPO-OE) or control (EV) conditions in three independent experiments (Figure S8A). At harvest, we confirmed complete BM donor reconstitution in all groups (Figure S8B). In the control conditions (EV-PBS and EV-PT), the blood parameters and BM morphology of EV-PBS and EV-PT mice did not differ (Figure S8C-E), indicating that PT treatment does not have major effects on the hematopoietic system, as shown previously (33).

In fibrosis, PT treatment also did not have major effects on myeloproliferation although there was a trend of lower platelets and WBCs (Figure S8C). The PT-treated TPO-OE mice had also a trend of higher BM cellularity (Figure S8D) and a trend of smaller spleens (Figure S8F). Strikingly, the reticulin fibrosis grade was significantly reduced in PT-treated compared to PBS control TPO-OE mice (Figure 8A). Furthermore, the spleen morphology of PT-treated TPO-OE mice was less disturbed, indicating less progressed MPN disease (Figure S8G). Importantly, we did not find major changes in HSC populations upon PT treatment, neither in EV nor in TPO-OE conditions (Figure S8H).

**Figure 8.**
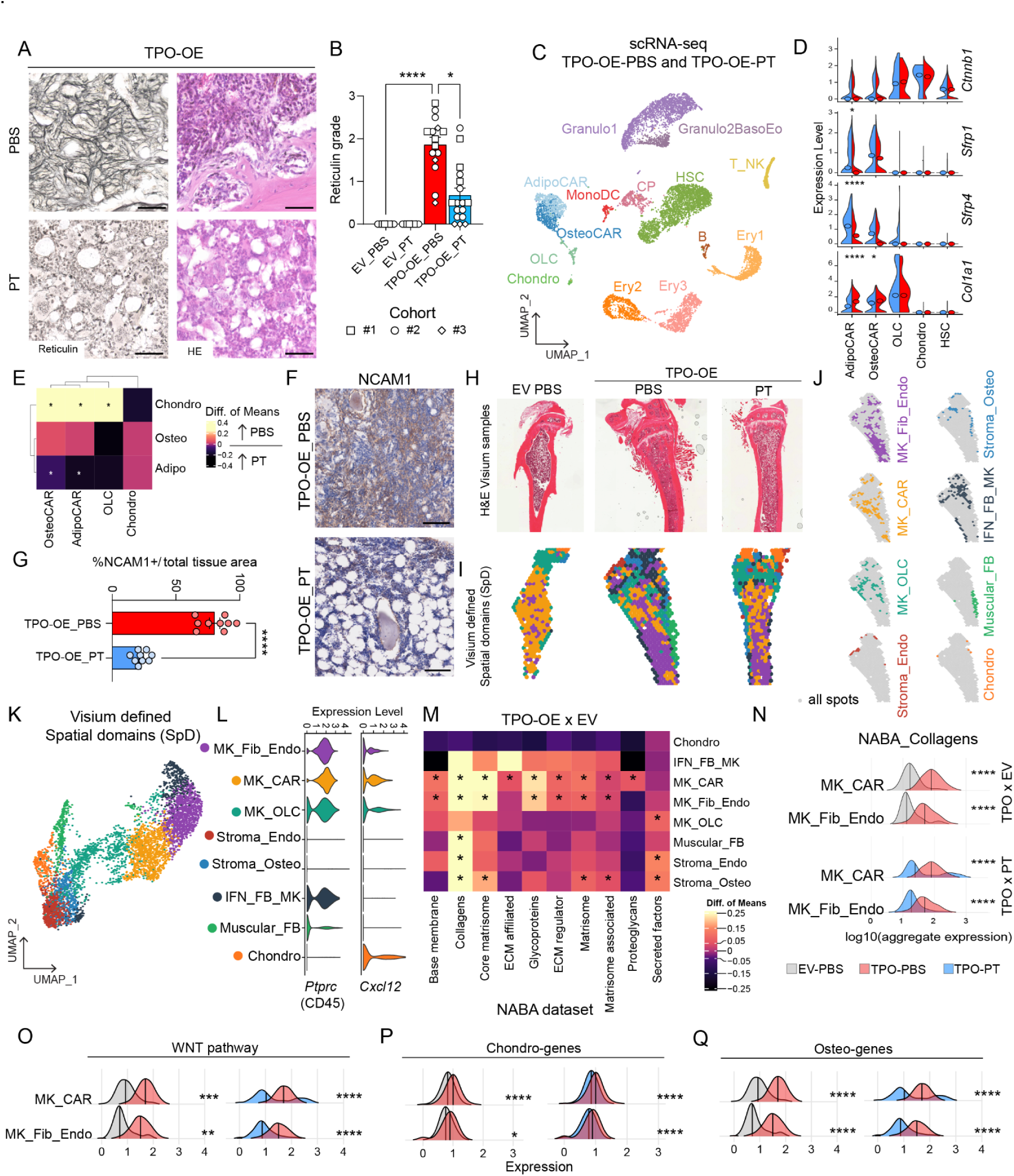
Inhibition of Wnt signaling reduces fibrotic transformation and osteoclerosis in BM fibrosis. (A) Representative Reticulin and H&E staining of tibias of BM transplant of cKit-enriched cells transduced with EV-eGFP (EV) or TPO-OE-eGFP (TPO-OE) viral vector and treatment with Pyrvinium-Tosylate (PT)-DMSO solution (PT) or PBS-DMSO (PBS) as a control. Scale bar: 50µm (B) Reticulin grade according to WHO. Kruskal Wallis rank test. EV-PBS n = 8, EV-PT n=8, TPO-PBS n=15, TPO-PT n =17, 3 independent cohorts, shape of point refers to cohort. (C) UMAP of scRNAseq dataset of lineage-depleted BM and collagen-digested bone chip fractions (digBC) of TPO-PBS and TPO-PT mice, n=3 mice pooled per experimental group. N= 11,063 cells. (D) Violin plot depicting the Wnt modulator *Ctnnb1* and inhibitors *Sfrp1*, *Sfrp4,* and *Col1a1* expression in stromal populations and HSCs. Median depicted with circle, wilcox test for significance. One sided Wilcoxon test. (E) Heatmap of differences of mean expression of adipo-, osteo-, chondro- genesets per stromal cluster from panel (O), TPO-OE_PBS compared to TPO-OE_PT. One sided Wilcoxon test. (F) Representative immunohistochemistry of NCAM1 in femurs of TPO-PBS and TPO-PT treated mice, space bar = 100µm. (G) Quantification of NCAM1 staining shown in panel M, n=3 mice per experimental group, quantification of 3 areas per slide, unpaired t-test. (H) H&E of tibias used for Visium spatial transcriptomics, n=3 per group, one representative image shown per experimental group. (I) Visualization of spatial domains (SpDs) detected in tibias as defined in (E) . (J) Individual SpDs shown in respective cluster colors with TPO-PBS bone as representative example. (K) UMAP of spatial domain clusters (SpDs) identified by Visium spatial transcriptomics. (L) VlnPlots of *Cxcl12* and *Ptprc* (CD45) expression per SpD. (M) Heatmap of differences of mean expression of NABA genesets per identified SpD, TPO-PBS compared to EV-PBS. One sided wilcoxon test used. High difference of mean expression represents higher expression in TPO-PBS compared to EV-PBS and vice versa. (N) Ridge plot showing aggregate expression of NABA collagen genes per identified SpD. Two-sided Wilcox test performed per spatial domain cluster, per comparison. One sided wilcoxon test was used to access the statistical significance of the distribution differences. (O - Q) Ridge plot showing aggregate expression of Wnt signaling pathway genes (O), chondro-genes (P), osteo-genes (Q) per SpD. TPO-OE PBS vs EV PBS in the left panel, TPO-OE PT vs TPO-OE PT in the right panel. Two-sided Wilcox test performed per spatial domain cluster, per comparison. Rest of SpDs shown in supplementary figure 8J. Statistical significance is indicated by: *p<0.05, **p<0.01, ***p<0.001, ****p<0.0001.

To demonstrate that the effect of Wnt inhibition by PT treatment was mostly on CAR cells and to a lesser extent on HSCs, we performed scRNA sequencing of the bone marrow after PT treatment compared to the PBS control and enriched for HSPCs and stromal cells (Figure 8C). We detected 11,063 cells (average 26,662 genes per cell) including major hematopoietic cell types as well as stromal cells (AdipoCar, OsteoCar and OLCs), which we annotated by comparison with our tdTom^+^ stromal clusters (Figure S8I). *Col1a1* expression was significantly reduced in adipo- and osteo-CAR cells upon PT treatment (Figure 8D). The Wnt inhibitors *Sfrp1* and *4*, which we identified to be significantly reduced in fibrosis (compare Figure 7C) in adipo- and osteo-CAR cells, were significantly increased in these populations upon PT treatment but not in HSCs (Figure 8D, S8J). Comparing the mean expression of chondro-, adipo- and osteo-associated gene sets also confirmed the upregulation of chondro- and osteo genesets in TPO-OE-PBS and an up-regulation of the adipo-genesets upon PT treatment (Figure 8E). These results, i.e. increase of Wnt inhibitor genes and normalization of stromal skewing signatures in CAR cells, support the direct effect of the PT treatment in the stromal compartment. Concordantly, the frequency of NCAM1+ cells throughout the BM was significantly reduced in TPO-OE transplanted mice after PT treatment compared to PBS-treated controls (Figure 8F, G). NCAM1+ cells were solely found in their peritrabecular localization, indicative of less activation of this population upon PT treatment.

To gain a spatial view of transcriptional changes occuring in PT-treated mice, we performed spatial transcriptomic experiments in nine bones (3x EV_PBS, 3x TPO-OE_PBS, 3x TPO-OE_PT) using the Visium platform (Figure 8H-J). We captured a total of 5,250 spots with an average recovery of 2,644 genes per spot. Unsupervised clustering identified eight spatial domains (SpDs)(Figure 8K, L). The majority of the dataset was represented by SpDs consisting of a mixture of CD45+ (*Ptprc* expressing) hematopoietic cells and Cxcl12-expressing stromal cells (Figure 8K, L) within the BM. The megakaryocyte(MK)-fibroblast (Fib) -endothelial (endo) SpD and the MK_CAR SpD mapped to the central marrow, while the MK_OLC SpD mapped to the growth plate, trabecular region and cortical bone of the diaphysis. The Chondrocyte (chondro) SpD aligned with the growth plate and other stromal populations were scattered across the bone (Figure 8J). We observed particularly high expression of platelet factor 4 (*Pf4)*, commonly expressed in monocytes, macrophages and megakaryocytes, in a number of SpDs (Figure S8K).

The most significant ECM-associated transcriptional reprogramming occurred in the MK_CAR and MK_Fib_Endo SpDs in BM fibrosis, particularly with regards to NABA collagens (Figure 8M). Strikingly, PT treatment significantly decreased collagen expression in these SpDs compared to TPO vehicle treated animals (Figure 8N). In line with our previous findings, we found increased Wnt pathway gene expression in the MK_CAR and MK_Fib_Endo populations in TPO bones compared to controls (Figure 8O, S8L). Furthermore, these SpDs had an increased chondrogenic, as well as osteogenic signature in fibrosis (Figure 8P, K, S8L), highlighting the calcification and osteogenic differentiation stromal switch occuring in BM fibrosis. These were also significantly reduced upon PT treatment supporting the notion that PT treatment prevents the stromal skewing in fibrosis and normalizes the stromal differentiation (Figure 8O-Q), as also observed in the scRNA-seq data (Figure 8E). We thus identified in an unbiased fashion a disease- and fibrosis-inducing cell-specific pathway which can be targeted by repurposing an FDA approved drug with a known safety profile.

## Discussion

There is increasing evidence that fibrotic processes across different organs are a result of a dysregulated repair response following repeated tissue injury, most notably during chronic inflammatory disorders (2,3,18,34). BM fibrosis is the most extensive remodeling of the BM and is most commonly initiated by the presence of mutated hematopoietic cells in myeloproliferative neoplasms, resulting in a reactive, cytokine- and chemokine-driven microenvironment.

Previously, Gli1^+^Lepr^+^ adipogenic- and osteogenic-CAR cells were shown to be functionally reprogrammed into fibrosis-driving cells in the presence of a fibrosis-inducing hematopoietic clone (8,11,35,36). However, spatial information on how the distinct anatomical regions of the bone are differentially affected was missing as either spatial resolution was not available or only the central marrow was taken into account. By combining fate tracing using three mesenchymal Cre-reporters, whole mount bone imaging and single-cell RNA sequencing, we now demonstrate a well-orchestrated interplay between the central marrow (perivascular niche) and the endosteal niche with focus on trabecular bone in fibrosis with high resemblance to skeletal repair involving the Wnt-mediated activation of a peritrabecular mesenchymal progenitor cell.

Our data indicate that the injury stimulus predominantly occurs in the central, perivascular BM niche, in line with recent reports (8,11,35,36). The central (perivascular) niche harbors 85% of hematopoietic stem and progenitor cells, including the mutated or fibrosis-inducing clone (37). Perisinusoidal and periarterial CAR cells, which mapped back to the central niche of the diaphysis, were functionally reprogrammed, lost hematopoiesis-supporting capacity, and acquired a pro-inflammatory phenotype, as a result of their increased communication with fibrosis-inducing hematopoietic cells Importantly, CAR cells showed a distinct phenotypic switch while leaving their perivascular niche and invading the central marrow. We provide evidence that true CAR cells are progressively lost during the fibrotic transformation through the upregulation of apoptosis and that their frequency is reduced. The absence of CAR cells in the direct association with endothelial cells exposes the endothelium to the inflammatory environment. Endothelial cells (Endo-1) showed transcriptional changes consistent with endothelial-to-mesenchymal transition. Apelin is a marker for H-vessels which represent the endothelium-bone axis and maintains the cross-talk of the metaphyseal and central bone marrow niche (24). In the context of our data, the upregulation of Apelin can be considered to promote increased communication and changes in the quality of the endothelium which coordinate the injury response of the perivascular central niche to the endosteal niche.

Within the metaphyseal region, stromal cells in close association with trabecular bone are a reservoir for fibrosis-driving cells in BM fibrosis. Cells residing in the peritrabecular region expand in a stepwise fashion during fibrosis and have a higher colony-forming potential compared to diaphyseal cells, indicative of a progenitor cell state. We propose that a subset of osteolineage cells, specifically OLC3, is activated in an injury-specific fashion and acts as an mesenchymal progenitor and is enriched in the Gli1-Cre driver in fibrosis. This was of particular interest as Gli1+ cells were previously identified as an osteogenic progenitor for fracture repair and termed “metaphyseal mesenchymal progenitors”. This postnatal Gli1+ cell pool with multi-lineage potential, immediately beneath the growth plate is essential for skeletal repair, but remains mostly quiescent in the adult mouse (12); (5). Importantly, we previously demonstrated that genetic ablation of Gli1+ cells rescues the fibrotic transformation of the bone marrow, proving their functional relevance in fibrosis (5). Our data now indicate that OLCs are skewed towards chondrogenesis and ossification (leading to osteosclerosis) and show increased differentiation into fibroblasts which are expanded in BM fibrosis. Ly6a+ fibroblasts express high levels of collagens at baseline and acquire an inflammatory phenotype, consistent with their behavior as first responders to injury in solid organs. So far, fibroblasts were under-represented and poorly described in bone marrow fibrosis single cell data sets, most likely due to the focus of previous reports on the central marrow (8,35). We identified NCAM1 as a marker for OLCs, which are expanded in fibrotic marrow, and correlate with fibrosis severity in primary myelofibrosis patients.

Functional differences between the perivascular and endosteal niche were also described during normal hematopoiesis, quite in line with our findings. The perivascular (sinusoidal) niche mostly reacts to injury (38), whilst the endosteal niche is important in regeneration. We demonstrate that mesenchymal progenitor cells with colony-forming potential are enriched in the metaphysis (peritrabecular region), in line with previous reports that perisinusoidal CAR cells are rather quiescent but have abundant cytokine expression (31) and that metaphyseal mesenchymal cells have self-renewing and multi-lineage differentiation potential (25). Peritrabecular regions are thus a hot-spot of fibrosis-formation and are activated in an injury-like fashion.

Based on our findings, the optimal anti-fibrotic treatment in the BM would target not only processes in the central marrow but also the peritrabecular progenitor cell. Receptor-ligand interactions demonstrated that OLCs are tightly regulated by Wnt signaling, in line with previous reports showing Wnt-mediated osteogenic transformation of bone marrow stromal progenitors in skeletal regeneration (31). Injury-induced Wnt signaling can drive mesenchymal progenitor cells into a reticular-osteoblast hybrid state and b-Catenin signaling was shown to be a driver but also therapeutic target in solid organ fibrosis (32), thus linking fibrosis and osteogenesis upon injury. Our data show that Wnt signaling is also upregulated in the myeloproliferative (non-fibrotic) phase in hematopoietic cells but fibrosis specifically correlates with stromal expression of b-Catenin in a disease-specific fashion. This is in line with recent studies identifying a TGFβ-Wnt-HOXB7 axis as associated with a pro-fibrotic and pro-osteoblastic biased differentiation of mesenchymal stromal cells isolated from patients, making this pathway an attractive target (39). Importantly, Wnt-mediated stromal cellular communications were mostly active in disease but not under steady state conditions, highlighting the potential to specifically target the diseased cells. Repurposing of Pyrvinium-Tosylate (PT) as a potentially clinically relevant compound that acts as an activator of casein-kinase 1a (CK1a or CSNK1a1) had anti-fibrotic effects and abolished osteosclerosis, restored the skewed differentiation of stromal cells in fibrosis, leading to the preservation of hematopoiesis. We demonstrated by scRNA sequencing that PT treatment affects predominantly (non canonical) Wnt signaling in stromal populations. Previous studies have, however, also shown a positive effect of MPN in general with other Wnt inhibitors, mostly reducing platelet counts (40), suggesting that targeting Wnt has positive effects on myeloproliferation but also on the fibrotic reprogramming of the stroma. We thus identified in an unbiased fashion a disease-, osteosclerosis- and fibrosis-inducing cell-specific pathway which can be targeted by repurposing an FDA approved drug with a known safety profile.

## Materials and methods

### Animal studies

All mouse studies were conducted according to protocols approved by the Central Animal Committee (Centrale Commissie Dierproeven [CCD], Netherlands) in accordance with legislation in The Netherlands (Approval No. AVD1010020173387 and AVD2216373). Mice were maintained on a 12-hr light/dark cycle and were provided with water and standard mouse chow *ad libitum*. Mice were randomly assigned to experimental groups.

PtprcaPepcb/BoyCrl (B6.SJL) and C57BL/6J mice were purchased from Charles River (Netherlands) and maintained in specific-pathogen-free conditions. Gli1CreERt2 (Gli1tm3(re/ERT2)Ali/J, JAX Stock #007913), Pdgfrb-creERt2 (B6-Cg-Gt(Pdgfrb-cre/ERT2)6096Rha/J, Rosa26tdTomato (B6-Cg-Gt(ROSA)26Sorttm(CAG-tdTomato)Hze/J, JAX Stock #007909), Ly6a-eGFP (B6.Cg-Tg(Ly6a-EGFP)G5Dzk/J, JAX stock #012643) and Grem1-creERT (B6.Cg-Tg(Grem1-cre/ERT)3Tcw/J, JAX stock #027039) were purchased from Jackson Laboratories (Bar Harbor, ME, USA). Offsprings were genotyped by PCR according to the protocol from Jackson laboratory. For lineage tracing, 10-15 week old mice were injected intraperitoneally with 4x 3mg tamoxifen in corn oil/3% ethanol (Sigma) at least 21 days before bone marrow transplantation. Unless otherwise specified, both male and female mice were used.

### Viral transduction for bone marrow transplant

Lentiviral particles were produced by transient transfection with lentiviral plasmid together with pSPAX and VSVG packaging plasmids using TranIT (Mirus) and concentrated by ultracentrifugation at 4°C for 2 hours. For lentiviral transduction, CD117(c-Kit)-enriched cells from donor mice were isolated by crushing hind legs and subsequent CD117-enrichtment by magnetic separation (Miltenyi Biotec). c-kit+ BM cells were cultured in StemSpan media (Stem Cell Technologies) supplemented by murine stem-cell factor (m-Scf, 50 ng/ml, Peprotech) and murine thrombopoietin (m-Tpo, 50 ng/ml, Peprotech). c-kit+ cells were transduced with virus (empty vector, EV-eGFP or thrompoietin overexpression, TPO-eGFP) in the presence of polybrene (4 ug/ml) for 48 hours, then transplanted into recipient mice in sterile saline solution.

### Induction of myelofibrosis by overexpression of Thrombopoietin (TPO)

Tamoxifen-induced recipient mice were irradiated with a split-dose regimen (2 x 6.02 Gy) and received cKit+ enriched cells from non-tamoxifen-treated littermates harvested 48h prior to transplantation and transduced with thrombopoietin overexpressing (TPO-eGFP+) lentivirus or control empty vector lentivirus (EV-eGFP+). For the Ly6a-EGFP and WT recipient cohorts, wild-type (WT) cKit+ enriched cells were transduced with thrombopoietin (TPO) or empty vector (EV) control lentivirus and transplanted into lethally irradiated (split-dose) B6.Cg-Tg(Ly6a-EGFP)G5Dzk/J (n= 5 mice per group) and B6.SJL recipient mice (n = 3-6 mice per group). Transplanted mice received drinking water supplemented with enrofloxacin (Baytril) for 3 weeks post-transplantation.

Blood was periodically collected *via* submandibular bleeds into microtainer tubes coated with K_2_EDTA (Becton Dickinson, NJ, USA) and complete blood counts were performed on a Horiba SciI Vet abc Plus hematology system. Mice were harvested when they had signs of fibrosis as indicated by the dropping of hemoglobin levels or weight loss according to defined humane endpoints. At harvest, long bones, pelvis and spine were collected for bone marrow analysis. Femurs were collected for imaging by fixing in 2% PFA for 10 hours, storage in 30% sucrose/PBS and processed further as described above/below.

### *In vivo* treatment with pyrvinium tosylate (PT)

Pyrvinium tosylate (PT) salt (SY-Pyrvinium) was purchased from Symansis (Timaru, New Zealand). The PT salt was first diluted in dimethyl sulfoxide (DMSO) at 10 mg/ml and stored at -80 C in dark tubes, and subsequently diluted in PBS for i.p. injection twice weekly at a dose of 0.5 mg/kg (PT groups). Control mice receive equivalent dilutions of DMSO in PBS (PBS groups).

### Histological and immunohistological analysis

Murine organs were fixed in 2% paraformaldehyde for 12h and transferred to 70% ethanol. Spleens were weighed before fixation. Bones were decalcified in 0.5M EDTA for 6 days, dehydrated, and paraffin embedded. H&E and reticulin staining were performed on 4 μm sections according to established routine protocols.

Human bone marrow biopsies were fixed for 24 h using the Hannover Solution (12% buffered formaldehyde plus 64% methanol), decalcified (EDTA), dehydrated and embedded in paraffin. Immunohistochemical stainings for active b-catenin (05-665, clone 8E7, Sigma) were performed with an automated, validated, and accredited staining system (Ventana Benchmark ULTRA, Ventana Medical Systems, Tucson, AZ, USA) using optiview universal DAB detection Kit (#760-700). In brief, following deparaffinization and heat-induced antigen retrieval the tissue samples were incubated with mouse Anti-Active-β-Catenin for 16 min at 36°C. Incubation was followed by hematoxylin II counterstain for 12 min and then a blue coloring reagent for 8 min according to the manufacturer’s instructions (Ventana). Slides were scanned and digitized in an automated fashion using a Hamamatsu Nanozoomer 2.0 HT system. Images were analyzed and exported using the NDP.view software (Hamamatsu, V2.5.19).

### Processing and whole-mount imaging of bone

Methods for 3D imaging of BM were adapted from previously published protocols (41). Mouse femurs were isolated, cleaned, and immersed in PBS/2% paraformaldehyde for 6 h, followed by a dehydration step in 30% sucrose for 72 h at 4°C. Femurs were then embedded in cryopreserving medium (OCT) and snap frozen in liquid nitrogen. Bone specimens were iteratively sectioned using a cryostat until the BM cavity was fully exposed along the longitudinal axis. The OCT block containing the bone was then reversed and the procedure was repeated on the opposite face until a thick bone slice with bilaterally and evenly exposed BM content was obtained. Once BM slices were generated, the remaining OCT medium was removed by incubation and washing of the bone slices in PBS three times for 5 min. For immunostaining, slices were incubated in blocking solution (0.2% Triton X-100, 1% BSA, and 10% donkey serum in PBS) overnight at 4°C. Primary antibody immunostainings were performed in blocking solution for 3 d at 4°C, followed by overnight washing with 0.2% Triton X-100 in PBS. Secondary antibody staining was performed for another 3 d at 4°C in blocking solution but in the absence of BSA to avoid cross-absorption. Immunostained thick femoral slices were successively washed overnight with 0.2% Triton X-100 in PBS and incubated in RapiClear 1.52 overnight. For observation under the confocal microscope, BM slices were mounted on glass slides while embedded in RapiClear. Confocal microscopy was performed with 10× (HCX PL FLUOTAR), 20× (HC PL APO CS2). and 63× (HCX PL APO CS2) in Leica STELLARIS 5 equipped with a 405-nm laser, white light laser and hybrid detectors. Confocal image stacks were processed and rendered into three-dimensional-volume on Imaris Software (Oxford Instruments, UK).

### Isolation of BM stromal cells for scRNA sequencing

Femurs, tibiae, hips, and spines were dissected and cleaned of surrounding tissue as much as possible. The bones were crushed in 2% FCS/PBS using a mortar and pestle. Dissociated cells were filtered through a 70μm nylon cell sieve, spun down and lysed with PharmLyse RBC lysis buffer. The cells were lineage depleted using biotinylated antibodies directed against lineages (CD5, CD45R, CD11b, Gr1, 7-4, Ter119) (Miltenyi Biotec) and additionally added CD45- and CD71-biotin antibodies (Biolegend). After staining for 10 min at 4°C, the cells were washed and incubated with anti-biotin beads (Miltenyi biotec) for 15 min at 4°C prior to magnetic depletion using a MACS column (BD).

For the isolation of cells from bones, crushed bone chips were washed with 2% FCS/PBS, and incubated with 10ml of collagenase II (1 mg/ml) at 37°C for 45 min under gentle agitation. The cell suspension was strained through a 70μm nylon cell sieve, spun down and lysed with PharmLyse RBC lysis buffer, before pooling with the lineage-depleted bone marrow fraction described above.

### FACS-staining and sorting of bone marrow stromal cells for scRNA sequencing

Cells were resuspended in 300μl PBS/2% FCS and stained at 4°C for 20 min with the antibodies described below. Washing was performed by adding 1ml PBS/2% FCS and centrifuging for 5 min at 300 × g, 4°C. After resuspension and addition of Hoechst (1:10,000), lineage/CD45/CD31 negative, tdTom positive cells were sorted into 400μl PBS/10% FCS (BD Aria III) and used as input for the 10x platform. A fully-stained sample from a tdTom negative mouse served as a negative control to define tdTomato gating. All antibodies were acquired from Biolegend.The following fluorochrome conjugated antibodies were used for murine samples: CD41-APC-Cy7, CD3-APC-Cy7, CD11b-APC-Cy7, Gr1-APC-Cy7, Ter119-APC-Cy7, B220-APC-Cy7, CD45.1-APC-Cy7, CD45.2-APC-Cy7, CD31-APC.

For non-tdTomato samples, TPO_PBS versus TPO_PT datasets, the same tissue processing as described above was performed, after which the sample was sorted for stromal (lineage/CD31/cKit/CD45/GFP negative alive singlets), and LSK events (lineage negative cKit+Sca1+ alive singlets). The stromal and LSK fractions were pooled in a 5:1 ratio and loaded onto the 10X chip. All antibodies were acquired from Biolegend.The following fluorochrome conjugated antibodies were used for murine samples: CD41-APC-Cy7, CD3-APC-Cy7, CD11b-APC-Cy7, Gr1-APC-Cy7, Ter119-APC-Cy7, B220-APC-Cy7, CD71-APC-Cy7, Sca1-PerCP, cKit-PeCy7, CD45.1-FITC, CD45.2-FITC, CD31-PE-CF594.

### Methocult (CFU-assay) of LSK sorted cells after PT treatment in vivo

1,500 primary, prospectively sorted LSK cells (lineage negative cKit+Sca1+ alive singlets) were plated out per 1ml Methocult GF M3434 +1%P/S in triplicate, with three biological replicates per experimental group. Plates were counted after 7 days.

### Culturing of bone marrow-derived stromal cells (cBMSCs)

Digested bone chips were washed thoroughly to remove hematopoietic contamination and cultured in full medium (alpha-MEM (Lonza), 20% FCS, 1% Penicillin/Streptomycin, 1ng/ml murine bFGF (Peprotech), 5ng/ml murine EGF (Peprotech) under normoxic conditions. After 9 days, the bone chips were discarded. Attached cBMSCs were passaged every 3-5 days at approximately 80% confluence and CD45/CD11b expression was assessed before start of experiment.

### Prospective sorting of stromal cells for CFU-F assays from metaphysis and diaphysis of long bones

Femurs, tibias and humeri were cleaned thoroughly, removing all muscle and tendons, and the periosteum was scraped using a small scalpel blade to remove potentially contaminating periosteal MSCs. The intact long bones were digested using trypsin (2.5mg/ml) and collagenase II (2mg/ml) for 20 minutes at 37°C, rinsed with PBS and scraped to remove any remaining periosteum. Long bones were cut into metaphysis and diaphysis and subsequently processed as separate samples (Meta/Dia). The bones were crushed in 2% FCS/PBS using a mortar and pestle. Dissociated cells were filtered through a 70μm nylon cell sieve, spun down and lysed with PharmLyse RBC lysis buffer. The bone chip fractions were digested with collagenase II (1mg/ml) for 45 minutes at 37°C then washed with 2%FCS/PBS, before combining with the respective BM fractions. Cells were stained for flow cytometry using anti-CD29, anti-CD31, anti-Sca1, anti-CD200, anti-VCAM1, anti-CD45, anti-Ter119, anti-B220, anti-CD3, anti-CD11b, anti-Gr1 and Hoechst as a live/dead marker. PdgfrbCre- or Gli1Cre-lineage stromal cells were sorted on a FACSAriaIII as live, lineage-negative, CD31- tdTomato+ cells and seeded in MesenCult Expansion kit (mouse, Stem Cell Technologies) according to the manufacturer’s instructions.

### Single-cell library preparation and sequencing

The libraries were prepared using the Chromium Next GEM Single Cell 3’ Reagent Kits (v3.1): Chromium Next GEM Single Cell 3ʹ GEM Kit v3.1 (PN-1000123), Chromium Next GEM Chip G (PN 2000177), Chromium Next GEM Single Cell 3ʹ Library Kit v3.1 (PN-1000157) and Dual Index Kit TT Set A (PN-1000215), and following the Chromium Next GEM Single Cell 3’ Reagent Kits (v3.1) User Guide (manual part no. CG000315, Rev B). Finalized libraries were sequenced on a Novaseq6000 platform (Illumina), aiming for a minimum of 50,000 reads/cell using the 10X Genomics recommended number of cycles (28-8-0-91 cycles).

### scRNA-seq data analysis

The scRNA-seq count matrix was obtained by aligning the raw sequencing reads into the custom td-tomato transcripts added mm10 mouse reference genome via the cellranger (version 4.0.0 and 6.0.1). Next, we used Seurat (v4.2) to analyze the scRNA-seq (42) based on R version 4.2.1. We filtered out cells with a high amount of mitochondrial genes (>10%), cells with low (<200) or high feature counts (>4,000), and removed the *Xist* gene to mitigate gender differences in downstream analyses. Next, we regressed out cell-cycle related genes, the proportion of mitochondrial, ribosomal, and UMI counts and performed a log-normalization of read counts. All scRNAseq libraries were integrated with Seurat’s reciprocal PCA (RPCA) based on the first 30 PCs. Unsupervised clustering was performed with the Louvain resolution 1.4. We used the FindMarkers gene function with an adjusted p-value < 0.05 to find cluster-specific makers. Differential gene expression analysis was performed, changing between distinct phenotypes. For this, we considered only genes with an absolute fold change greater than 0.5 and adjusted p-value < 0.1. GO and pathway enrichment analysis was based on the EnrichR package looking into the NABA gene set (21). All p values were corrected by Benjamini-Hochberg correction. Differentially expressed genes, including corresponding statistics are supplied in a table inside the folder for Figure 2 of the 08-11-2024 Zenodo repository.

scRNAseq analysis, histological stainings and microCT analysis are described in detail in the supplemental materials.

## List of Supplementary Materials

### Tables

**Table S1** List of differentially expressed (DE) genes between TPO-OE and EV, related to figure
2

**Table S2** Gene sets related to Figures 5 and 8

**Table S3** Human cohort characteristics related to Figure 6

## Acknowledgement

R.K.S. is an Oncode Investigator and is supported by ERC grants (Rewind-MF ERC-CoG 101124542; deFIBER ERC-StG 757339 and PoC DeAlarmin) and a ZonMW VIDI grant. This work was in part supported by grants of the Deutsche Forschungsgemeinschaft (DFG) (German Research Foundation) to H.G (417911533), R.K. (KR 4073/9-1), R.K.S. (504777725; 417911533; 514007497) and I.C. (417911533). The project received funding from the program “Netzwerke 2021”, an initiative of the Ministry of Culture and Science of the State of Northrhine Westphalia (CANTAR network). R.K., I.G.C., and R.K.S. are members of the E:MED Consortia Fibromap and the consortium CureFib funded by the German Ministry of Education and Science (BMBF). H.G. is supported by a Gilead Research Scholar Award in Oncology/Hematology, a ZonMW VENI grant and an Erasmus Medical Center Fellowship.

We thank Remco Hoogenboezem for pre-processing of the scRNA sequencing data; Stephani Schmitz, Aurélie van Kleef-Boeree, Iris Bakker and Pia Wanner for technical support; Juliette Pearce for drawing the graphical abstract art; the Developmental Biology department of Erasmus MC for helpful scientific discussions.

## Author contributions

Conceptualization: RKS, IGC, RK, CNA, BB, JN

Methodology: RKS, IGC, RK, CNA, BB, JN

Investigation: BB, JN, HG, YM, IS, SF, JB, JP, US, ARM, KKS, RHA, JG, RK, RKS

Whole-mount imaging: YM, CNA

Quantification tdTomato signal: HTM

Image analysis and quantifications IHC: AKG

Visium spatial sample preparation: AB

μCT analysis: MR, MW, RBS, TL, FK

Patient tissue collection, biobanking, analysis human biopsies: HR, NKA, DR

MarrowQuant analysis: RS, ON

Sequencing: EMJB

Visualization: RKS, IGC, HG, BB, JN

Funding acquisition: RKS, IGC

Project administration: RKS, IGC

Supervision: RKS, IGC

Writing – original draft: BB, JN, RK, CNA, IGC, RKS

Writing – review & editing: BB, JN, HG, IGC, RKS

## Supplemental Materials and Methods

### Single-Cell Proportion Analysis

Differential Proportion Analysis was performed using the package scProportion (Version 0.0.0.9; https://github.com/rpolicastro/scProportionTest), accessing the cell abundance difference in the phenotypes presented in the study.

### scRNAseq niche localization prediction with cell deconvolution and label transfer

We made use of laser capture data (LCM-seq) composed of five distinct bone marrow niches in homeostasis (Arteries, SubEndosteum, Endosteum, High sinusoids, and Low sinusoids) to recover the spatial context from the scRNA-seq data(16), samples from sinusoid sections were disregarded due to low mapping of signatures as the sinusoidal bulkRNAseq contained mainly hematopoietic cells. We defined a gene expression signature for each niche by using the ROC statistics from Seurat. We considered all genes presenting a ROC value lower than 0.3 or larger than 0.7. We then used these signatures as reference for deconvolution analysis by providing single-cell data from Homeostasis samples as input with a CIBERSORTx algorithm(43) implementation provided in (https://github.com/veltenlab/rnamagnet,(16)). We summed up the probabilities for all cells in a cluster, providing a niche score for each cell type. For recovery of metaphyseal and diaphyseal location signatures, we made use of TransferLabels in Seurat, using a published scRNA-seq dataset containing both metaphyseal and diaphyseal BM stromal populations (15).

### Osteogenesis, Chondrogenesis and Adipogenesis genesets

The osteo/adipo genesets were obtained from Supplemental Figure S7(25) and chondro geneset from Supplementary Table S2(27). The median of the genesets gene expression was compared using the Wilcoxon Test. We have added these datasets in Table S1.

### Coefficient of variation (CV) scoring - SPEC (differentiated) and MONO (progenitor) scores

We explored the coefficient of variation (CV) score to measure the stem-ness or specialization state of a cell. For every gene, the CV is calculated as the standard deviation divided by average expression. Low CV is indicative of genes with low variance (i.e. not being cell specific), while high CV indicates cell specific genes (as cells start expressing specific genes once they decide on a fate). We next perform a z-score transformation of the gene CV scores. Genes with z-score below 0.5 are defined as low CV genes, while genes with z-score > 0.5 as high CV genes. Only genes expressed in more than 50 cells are considered. We finally obtain a CV-low score per cell as the total expression value of low-CV genes, and the CV-high score the total expression of high-CV genes per cell. High-CV scores (SPEC score) indicate the level of specialization of the cell, i.e. cells whose gene expression is dominated by cell specific genes.

Low-CV scores indicate cells with monotonous (or stem-like properties), as they express genes common to most cells (MONO score).

### Reconstructing cell development trajectories

For trajectory analysis, we used phateR implemented in the Seurat package (runPHATE) with k=20 and the four Phate components(44). Next, the Phate components were used to re-cluster the cells, and the trajectories were defined using the new clusters and the CV score described above. As described in the ArchR package, the trajectories were obtained by computing the smooth::spline with the N-dimensional coordinates and the pseudo-time values.

### Ligand-receptor (LR) analysis

For ligand-receptor analysis, we used the CellPhoneDB method and a LR consensus database implemented in the LIANApy package (Version 0.1.10,(45)). We performed the analysis per phenotype (Homeo, EV, and TPO). We only consider statistically significant LR pairs (p-value < 0.01). We next used CrossTalkeR(Version 1.3.2,(30)) to characterize major phenotype-specific cell-cell interactions. We only considered cell pairs, in which the number of LR pairs changes significantly (Fisher’s Test, p-value < 0.05) between two compared conditions. We used GO Biological Process 2011 database (available in https://maayanlab.cloud/Enrichr/#libraries) to subset pathway-specific (Wnt) LR interactions.

### Ligand-receptor (LR) category-based filtering

Cell communication scores obtained through CrossTalkeR(30) were subset according to the Ligand classification from the Matrissome database(46) or the Cytosig database(46), and the following cell niches:

Metaphysis = [“OLC1”, “OLC2”, “OLC3”, “Pre_chondro”, “Chondro”, “FB1”, “FB2”, “FB3”, “FB5”, “FB6”, “FB7”, “OB”]

Transition = [“Epi”, “FB4”, “IFN_FB”, “CAR7_ribo”, “Mus_CAR”, “FB8”, “Stroma”, “Endo2”, “Mural”, “Endo1”, “Smooth_Mus”]

Diaphysis = [“IFN_CAR”, “CAR1”, “CAR2”, “CAR4_Pdg”, “CAR3”, “CAR6”, “CAR5”]

Hematopoietic (Hemap) = [“Hemap_prog”, “Neutro1”, “Neutro2”].

The significance of the differences between the distributions of subset cell communication scores in different conditions (TPO and EV) was addressed through two-sided Wilcoxon tests as implemented in the Scipy library (version 1.9.1) in Python(47).

### Bone marrow computational pathology quantification MarrowQuant

The MarrowQuant (mouse version) algorithm was employed for bone marrow tissue quantifications. Within QuPath 0.3.2, annotations were performed following the methods described in previous studies (Sarkis et al., 2023; Tratwal et al., 2020). In brief, tissue boundaries encompassed most of the tissue, excluding highly hemorrhagic regions. The bone area was excluded from the analysis, and the total marrow area is defined as the selected region of interest minus the bone and artifact areas therefore defined as the denominator. The MarrowQuant algorithm was subsequently executed, with a minimum size threshold set at 120µm2 for adipocytes. The resulting quantification data were exported in a .txt file. Codes and tutorials are available on the following GitHub link: https://github.com/orgs/Naveiras-Lab/repositories

### StarDist on Adipocytes

For the individual segmentation of bone marrow adipocytes and tracking of their size distribution, we utilized the StarDist model trained specifically for bone marrow adipocytes, as detailed in Sarkis et al. 2023 (48). This algorithm was also integrated within QuPath, and the same tissue boundaries defined for MarrowQuant quantifications above were applied to StarDist. Consequently, StarDist accurately detected the individual adipocytes. Subsequently, a classification algorithm was employed, and the quantification results were exported using a .txt file. Codes and tutorials are available on the following GitHub link: https://github.com/orgs/Naveiras-Lab/repositories.

### microCT scans of tibia and analysis

A Bruker Skyscan 1272 microCT scanner equipped with a 11 megapixel sensor was used. The following parameters were chosen for scanning: 4x4 µm^2^ pixel size and a matrix of 1344x2016. The voltage was 60 kV and the current 166 µA. Furthermore, a 0.25 mm Al filter was applied. The exposure time per image was set to 1558 ms and 8 averages per projection were acquired. The step between projections was set to 0.2° and a total of 180° was imaged, resulting in 900 projections per scan. Data reconstruction and beam hardening corrections (up to 70%) were performed using the NRecon software (Bruker, Belgium). Morphometric and densitometric analysis were done in CTan software (Bruker, Belgium) inside a cylindrical VOI (7.07 mm^2^ x 1.6 mm) located in the upper part of tibia from the section where secondary spongiosa forms a sole uninterrupted region. A sequence of image processing steps was applied to virtually separate the trabeculae area from cortical bone. After bone thresholding, the outer surface of the tibia was defined by the ROI-shrink wrap method. Inside this ROI, the inverted binarized image was treated by a morphological escalation method (opening and closing with a stepwise increasing parameter from 2 up to 10px). After removal of residual speckles, the region of trabeculae was defined. Then, 3D morphological analyses were performed on the new binarized image inside this trabeculae VOI and following parameters were obtained: bone volume to total volume (BV/TV); trabecular number (Tr.N); trabecular thickness (Tr.Th) and connectivity of trabeculae (Connectivity).

### Quantification of tdTomato cells

All image stacks were processed using FIJI (49). Briefly, images (n=3 per condition) were contrast enhanced, normalized, and converted to 8-bit grayscale. Contrast enhancement and normalization were performed to standardize the intensity levels across all images, ensuring comparability. The intensity profile along multiple lines perpendicular to the bone surface was measured to ensure representative sampling across the bone surface. All profile line measurements were done from the start of the growth plate border, which corresponded to the highest recorded intensity values. The total integrated density along the distance for each condition was calculated by averaging the integrated density values of all lines across all images. The area under the curve (AUC) for each condition was calculated using the trapezoidal rule after the growth plate border. Statistical significance between conditions was determined using a t-test, with a p-value <0.05 considered significant.

### Quantification of NCAM1/CD56-stained IHC murine sections

All images were analyzed using ImageJ2 version 2.14.0. Briefly, three representative regions of the BM per sample were selected using the rectangle tool, and total tissue area was measured after color thresholding. Afterwards, we made use of the ‘color deconvolution’ plugin for ‘FastRed/FastBlue/DAB’ to retrieve the signal from the DAB channel (NCAM1/CD56). The threshold for the positive signal of DAB was determined using automatic thresholding schemes per image, and measured as the percentage area of the image.

### Visiopharm image analysis of NCAM1/CD56-stained human sections

The Visiopharm Integrator System (VIS) platform version 2023.01 was used to analyze digitized serial human BM slides. Image analysis protocols are implemented as Analysis Protocol Packages (APP) in VIS. Several APPs were designed to quantify slides stained with CD56. Prior to the image analysis, it was important to outline the Region of Interest (ROI), hence a number of auxiliary APPs were designed to detect ROIs. To detect ROIs, the first auxiliary APP ran on the slide using threshold classification to identify the tissue regions. The second auxiliary APP ran on the slide using the DeepLabv3 network of the VIS AI module to identify the bone and peeled regions of the biopsy. For the analysis, an APP ran on 20X on the slides using the U-Net network of the VIS AI module to identify NCAM1/CD56-positive staining. As a post-processing step, a threshold of 22 pixels away from bone or peeled areas was used to classify the detected NCAM1/CD56 positive staining into two classes and output variables obtained from the APPs include: total tissue area, total NCAM1/CD56 positive area, NCAM1/CD56 positive area close to bone or peeled areas and NCAM1/CD56 positive area away to bone or peeled areas. The CIF score was determined as previously described (29).

### Prospective sorting of NCAM1+ and VCAM1+ BM stromal cells

For the prospective sorting of NCAM1/CD56+ and VCAM1/CD106+ BMSCs, we processed samples as above in ‘Isolation of BM stromal cells for scRNA sequencing’, and sorted lineage/CD31 negative alive singlets NCAM1+ or VCAM1+. After three passages the cells were immortalized using Large-T virus and characterized using flow cytometry and RT-qPCR.

### NCAM1 and VCAM1 stromal cell co-culture with cKit cells

c- Kit+ hematopoietic stem and progenitor cells were isolated from WT mice using the autoMACS pro Separator after crushing the compact bone (Miltenyi Biotec). c-Kit+ cells were seeded on top of immortalized NCAM1+ or VCAM1+ cBMSCs in StemSpan™ Serum-Free Expansion Medium (Stem Cell Technology, Vancouver, Canada) supplemented with murine thrombopoietin (m-tpo) (50 ng/mL; Peprotech), murine stem cell factor (m-scf) (50 ng/mL; Peprotech). After 24 hours of co-culture, the non-adherent hematopoietic cells were collected for apoptosis flow cytometry (Annexin V APC Apoptosis Detection Kit, eBioscience). All samples were analyzed by flow cytometry using a FACS Fortessa (BD Biosciences, San Jose, CA). Data were analyzed using FlowJo software (Version 10, TreeStar Inc.)

#### RNA extraction and real-time qPCR analysis

RNA from cBMSCs was extracted using Trizol solution (ThermoFisher) according to the manufacturer’s instructions, and 1,0 μg of total RNA was reverse transcribed using the high-capacity cDNA Reverse Transcription kit (Applied Biosystems). Quantitative polymerase chain reactions were performed with SYBRGreen PCR master mix (ThermoFisher) on an Biorad CFX384 Real-Time PCR System. Glyceraldehyde-3-phosphate dehydrogenase (Gapdh) was used as a housekeeping gene. Data was analyzed using the -Δct method. Primers are listed in the table below.

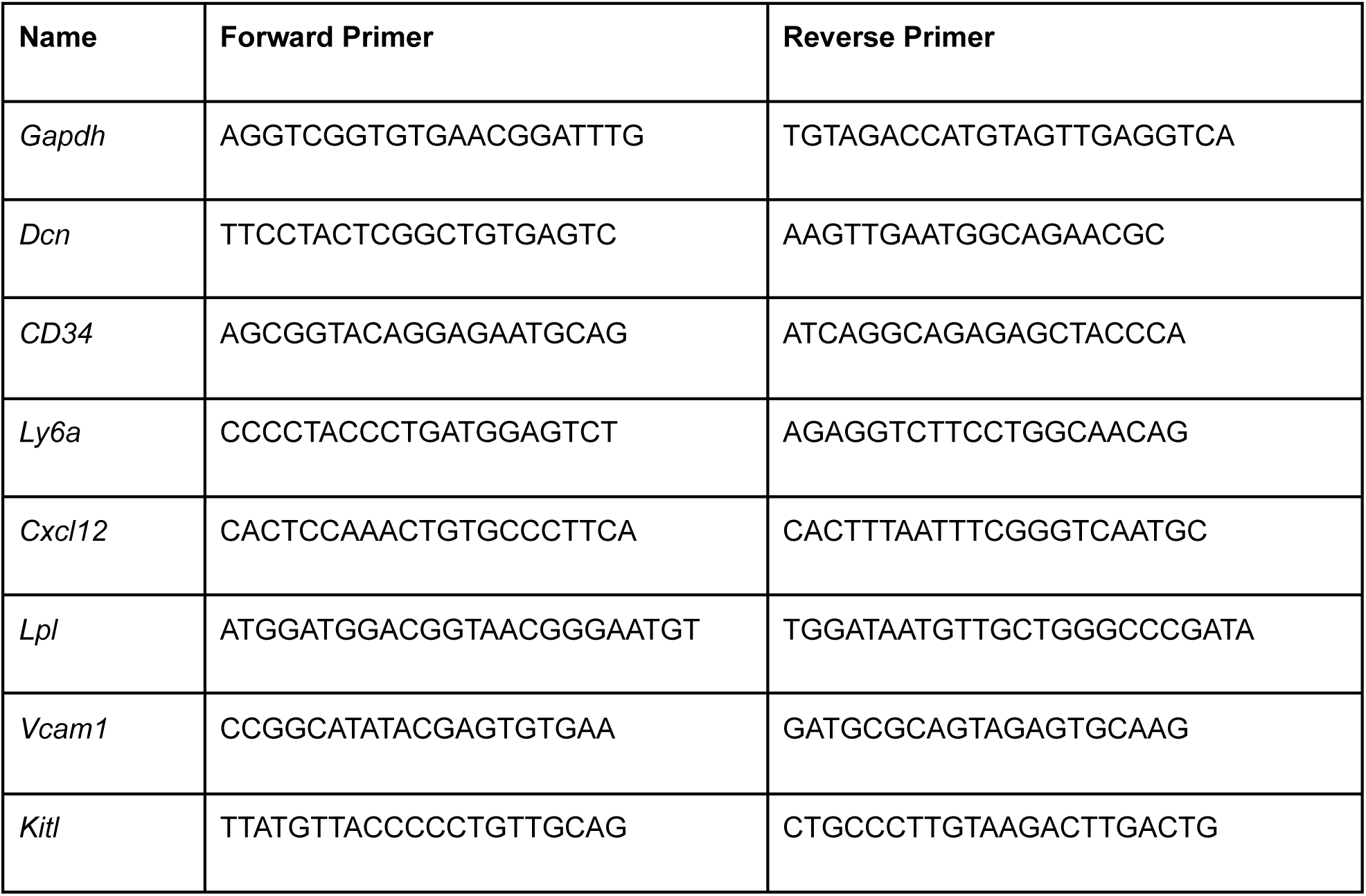
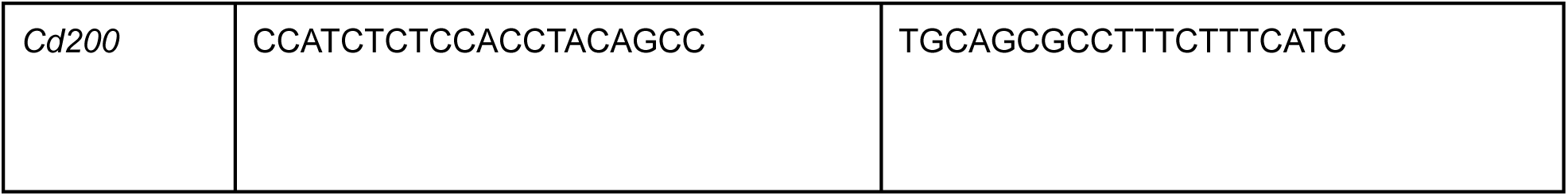

### Visium spatial transcriptomics sample preparation

Visium CytAssist Spatial Gene expression assay (10X Genomics, PN #1000523) was conducted following 10X Genomics recommendations. Briefly, three FFPE sections were placed onto each capture area of a 11mm2 Visium CytAssist slide (three experimental groups per slide, three replicates per experimental group). Following section placement, deparaffinization, H&E staining, brightfield imaging, decrosslinking and library preparation were performed. Libraries were sequenced on a NovaSeq sequencer (Illumina) using sequencing depth and parameters recommended by 10X Genomics.

### Visium spatial transcriptomics analysis

For each bone presented in each slide, a segmentation file was generated as recommended by 10x Genomics using Loupe Browser. Next, sequencing reads and the alignment files were processed using spaceranger v.3.0.0. Seurat was used to filter low quality cells (nFeature_Spatial>200 & nFeature_Spatial < 7500 & percent.mt < 5) per sample. Data integration was done using harmony and spatial domains (SpDs) were identified using the FindCluster with resolution=0.7.

### Single cell RNA sequencing of Pyrvinium Tosylate (PT) treatment cohort

The scRNA-seq count matrix was obtained by aligning the raw sequencing reads into the mm10 mouse reference genome via the cellranger (version 8.0.1). Seurat (version 5.0.1) was used to filter low quality cells (nFeature_Spatial>200 & nFeature_Spatial < 7500 & percent.mt < 5) per sample. Data integration was done using harmony and single cell clusters were identified using the FindCluster with leiden clustering using resolution=0.1. Stroma cells’ clusters were further sub-clustered using FindCluster with leiden clustering using resolution=0.2.

---

**Figure S1 (related to figure 1):**
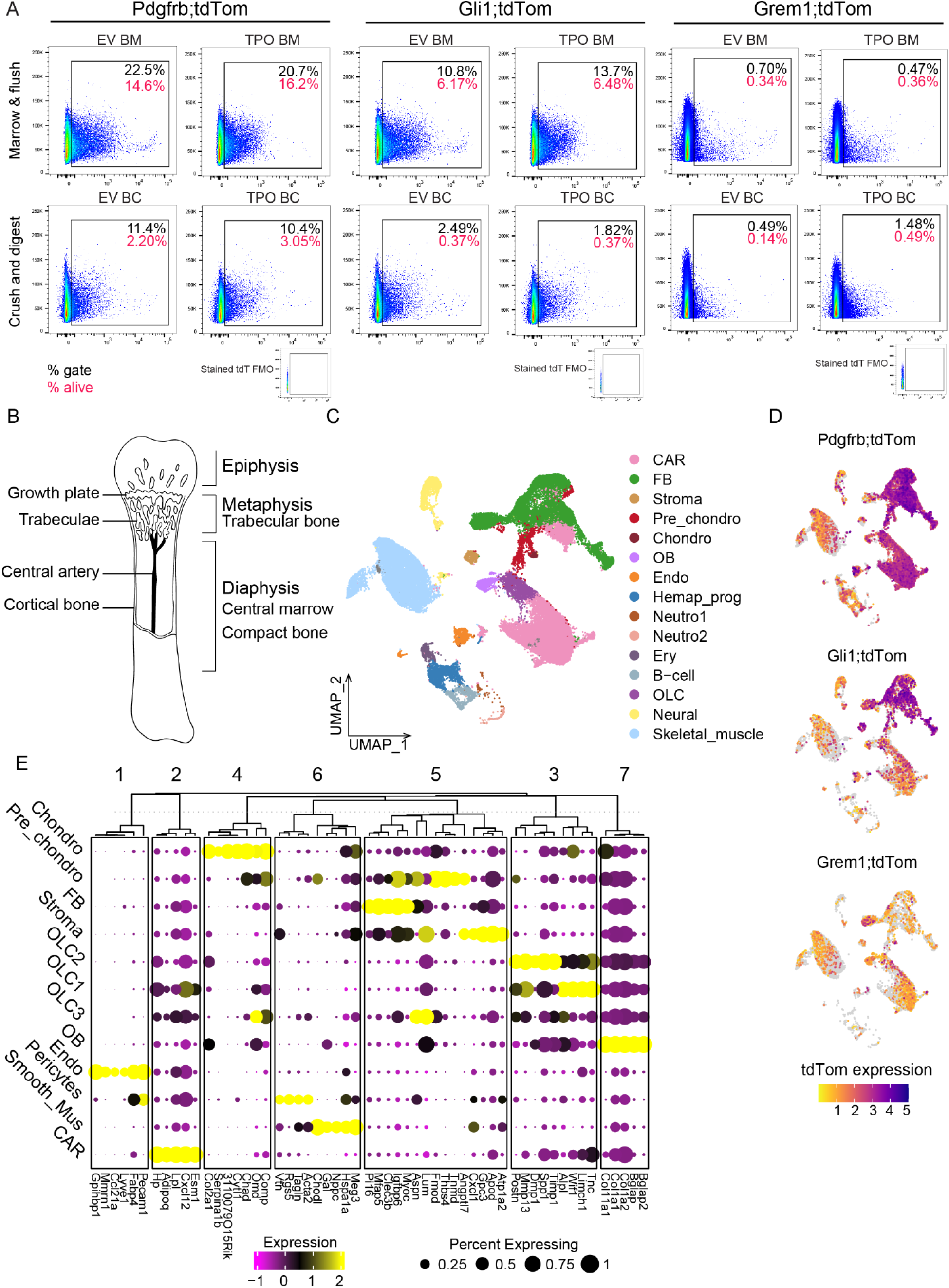
The combination of stromal Cre drivers provides high granularity of the bone marrow niche. (A) FACS plots showing tdTomato expression in the different fractions of the bone marrow (BM) per Cre-driver. EV= empty vector control; TPO=Thrombopoietin-induced bone marrow fibrosis (B) Schematic of anatomical structures and niches in the bone marrow (C) Whole dataset before removal of skeletal cells, hematopoietic cells (D) Expression of tdTomato transcript per Cre-reporter (E) Dot plot of top 5 markers per cluster of re-clustered stromal dataset

**Figure S2 (related to figure 2):**
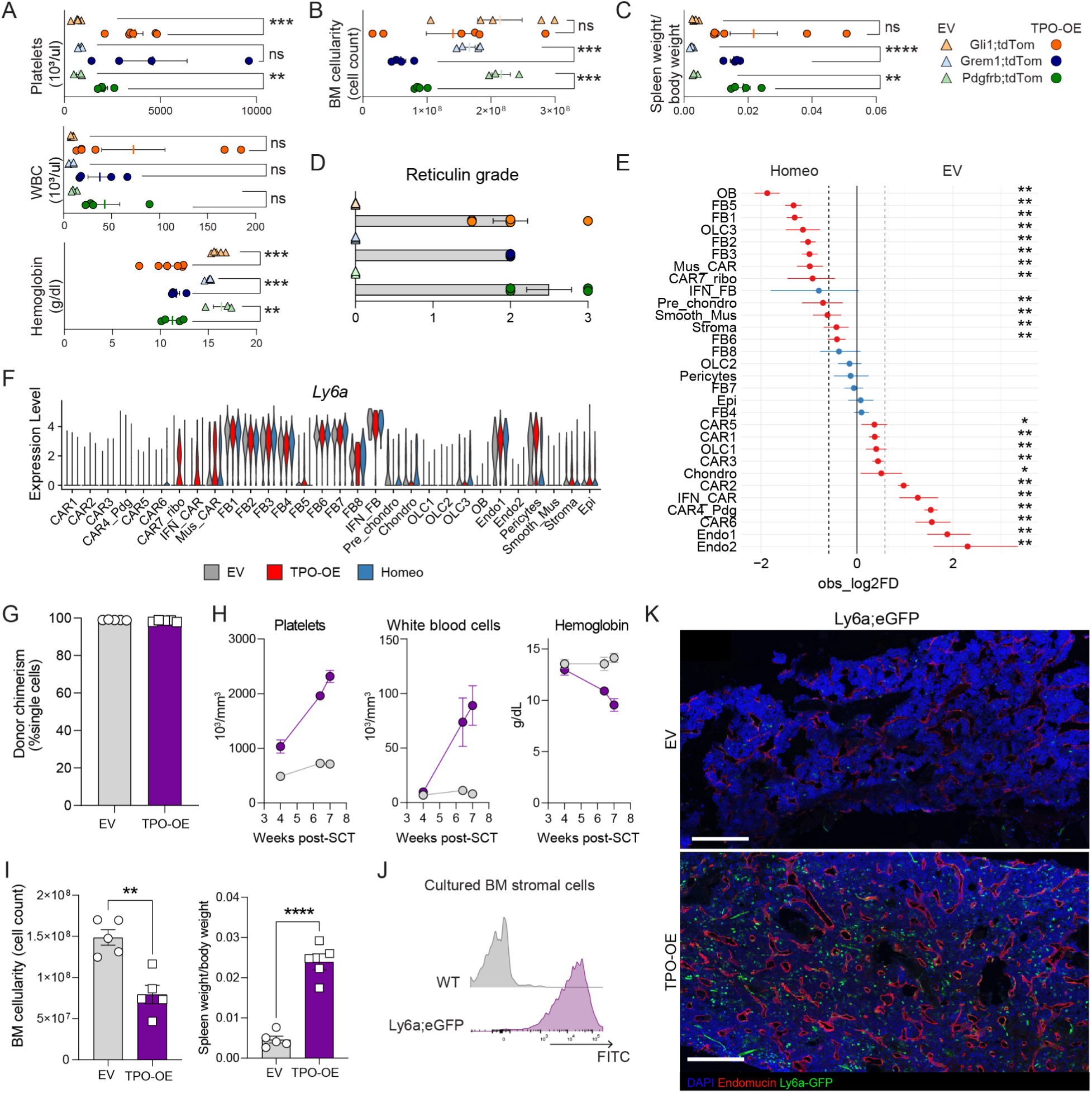
CAR cells acquire a pro-fibrotic phenotype but are reduced in frequency while fibroblasts expand in bone marrow fibrosis. For panels A, B, C and D: Gli1;tdTom: EV n=5, TPO-OE n = 6, Grem1;tdTom: EV n=4 TPO-OE n=4, Pdgfrb;tdTom: EV n=3, TPO-OE n=4. (A) Blood counts of experimental mice used for 10X scRNA sequencing, t-test performed per genotype group ( Pdgfrb;tdTom, Gli1;tdTom, Grem1;tdTom). (B) Cell counts of the bone marrow (BM cellularity) at the end of the experiment of 10X cohort; t-test performed per genotype group ( Pdgfrb;tdTom, Gli1;tdTom, Grem1;tdTom). (C) Spleen weight/body weight at the end of the experiment of 10X cohort. t-test performed per genotype group ( Pdgfrb;tdTom, Gli1;tdTom, Grem1;tdTom. (D) Fibrosis grading (reticulin grade) of BM of 10X cohort. t-test performed per genotype group ( Pdgfrb;tdTom, Gli1;tdTom, Grem1;tdTom). (E) Proportion changes in Homeo versus EV obtained using scProportionTest. (F) Violin Plot of *Ly6a* expression per cluster per condition (G) Donor chimerism after transplant, n=5 mice per group in the Ly6a;GFP recipient transplant; In brief, Ly6a;eGFP mice were lethally irradiated and intravenously received c-kit-enriched HSCs from WT littermates expressing either thrombopoietin cDNA (TPO-OE; n = 5, three males) or control cDNA [empty vector, EV, n = 5; both lentiviral SFFV-iblue fluorescent protein (BFP) vector backbone] as outlined in Figure 2A. (H) Hematological phenotype in Ly6a-eGFP transplanted animals comparing to the empty vector; (EV) control condition (grey) to Thrombopoietin-overexpression (TPO-OE; purple); n=5 mice per group. (I) BM cell counts (bone marrow cellularity) and spleen size expressed as spleen to body weight ratio. One-way ANOVA with multiple comparisons. (J) Stromal cells isolated from bone chips of Ly6a-GFP mice show strong GFP signal in culture as indicated by flow cytometry. (K) Representative images of EV control and TPO-OE BM of Ly6a;GFP mouse, Endomucin in red, DAPI nuclear stain in blue, scale bar = 100µm Statistical significance is indicated by: *p<0.05, **p<0.01, ***p<0.001, ****p<0.0001.

**Figure S3 (related to figure 3).**
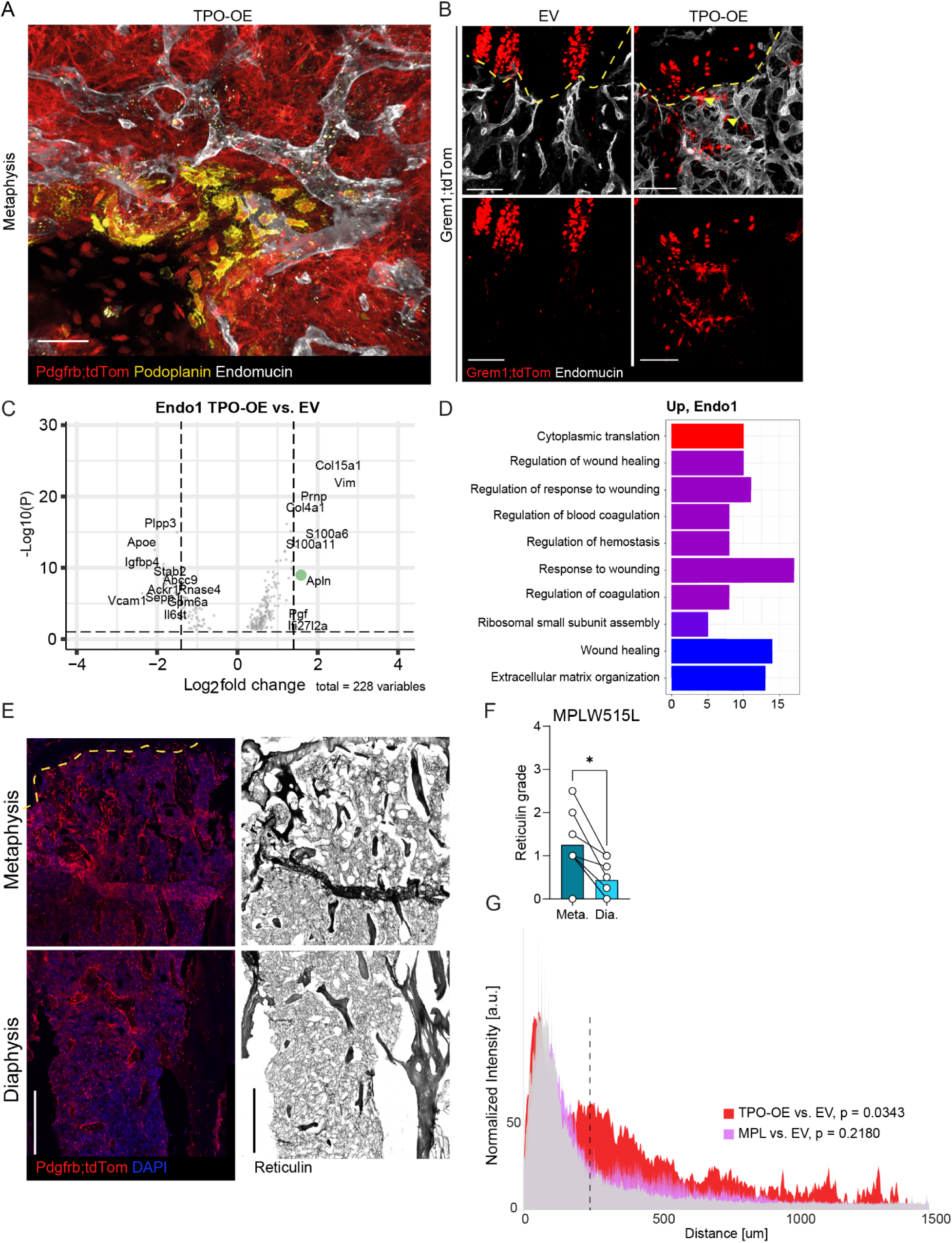
Distinct phenotype switch in TPO-OE induced bone marrow fibrosis in the perivascular (diaphysis) and endosteal (metaphysis) localization. (A) Zoom in of peri-trabecular region in TPO-OE (fibrosis) transplanted Pdgfrb;tdTom reporter mice showing region of osteosclerosis (new bone formation), marked with podoplanin staining and typical morphology of osteo-lineage cells/osteoblasts. Scale bar: 50µm. (B) Whole-mount imaging of Grem1;tdTom in the control (EV) and fibrosis (TPO-OE) condition. Reticular-like cells emerging from growth plate (GP) indicated with yellow arrow head. Dotted line = growth plate. Scale bar: 50µm (C) Volcano plot showing differentially expressed genes (TPO vs EV) in the endothelial cluster endo-1. Apelin (Apln) is highlighted in green. (D) Gene set enrichment of differentially expressed genes in endo-1. (E) Confocal images of fibrotic bigenic PdgfrbCreER;tdTomato depict the peritrabecular region as a hotspot zone in bone marrow fibrosis in particular in comparison to the central marrow (diaphyseal) region (zoom in). Demounted and reticulin stained whole mount bone (shown in E). The hotspot tdTomato-positive areas overlap with increased reticulin deposition (peritrabecular region/metaphysis), Representative regions of interest (ROIs) are shown along the tibia. Scale bar: 500µm. (F) Grading of reticulin fibrosis grade according to WHO criteria in the metaphysis (meta) and diaphysis (dia) in femurs of mice transplanted with ckit-enriched cells transduced with the MPLW515L mutation. (G) Quantification of Gli1;tdTom+ cell expansion/movement away from growth plate (GP) border in femurs of mice transplanted with ckit-enriched cells transduced with the MPLW515L mutation. Statistical test: two-sample t-test on area under curve (auc), with a p-value <0.05 considered significant.

**Figure S4 (related to figure 4).**
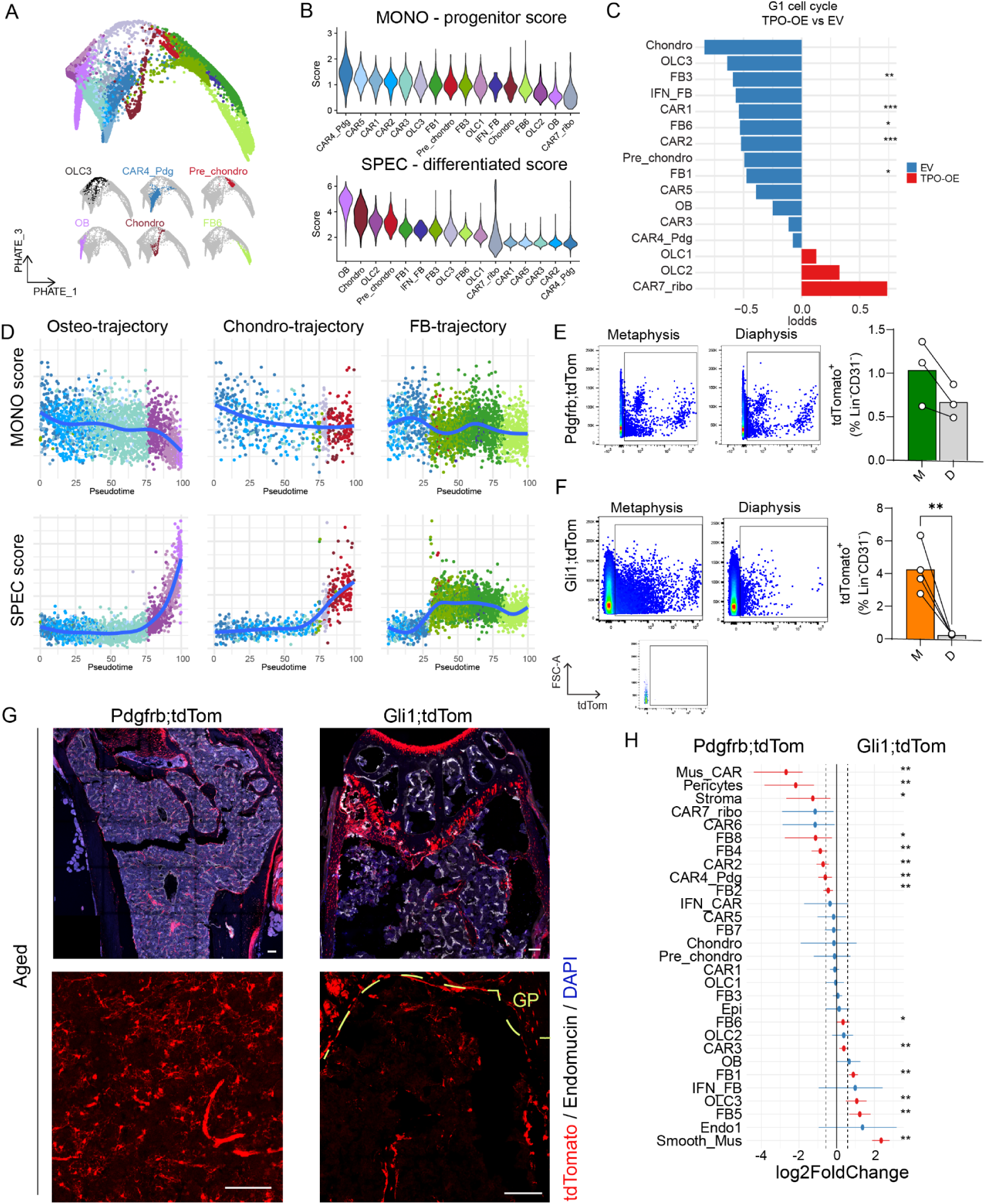
Stromal differentiation trajectories. (A) PHATE dimension reduction differentiation trajectory highlighting the distribution of the included clusters as highlighted in the lower panel. (B) Violin plots of the progenitor (MONO) and differentiated cell (SPEC) score per celltype. X-axis is ordered according to the mean values. (C) G1 cell cycle analysis of stromal cell clusters, comparing TPO-OE vs. EV control. Statistical comparison was done using Fisher Exact Test. (D) Progenitor (MONO) and differentiation (SPEC) scores along trajectories defined in Fig. 4C (E, F) Flow cytometry analysis of TdTomato expression in metaphyseal versus diaphyseal regions in Pdgfrb;tdTom (E) and Gli1;tdTom (F) homeostatic bones, in the right panel quantification of tdTom+ cells per region, paired t-test. M. = metaphysis, D. = diaphysis. Insert shows fluorescence minus one (FMO). (G) Representative confocal images of aged Pdgfrb;tdTom and Gli1;tdTom mice, 1 year after tamoxifen induction. Note the wide-spread abundance of Pdgfrb;tdTom-lineage cells (diaphysis, metaphyseal region), whereas Gli1;tdTom-lineage cells are located at the metaphyseal region and growth plate. Dotted yellow line represents growth plate (GP) (H) Proportion test Gli1;tdTom versus Pdgfrb;tdTom, on homeostasis subsetted data, obtained using scProportion test. Relative differences in cell proportion per cluster, red colored dots show significant fold change (FDR<0.05 and absolute fold change > 0.58) with error bars showing confidence intervals for the magnitude difference (permutation test, n=1000). Blue dots- ns; red dots: significant changes. Statistical significance is indicated by: *p<0.05, **p<0.01, ***p<0.001, ****p<0.0001

**Figure S5 (related to figure 5):**
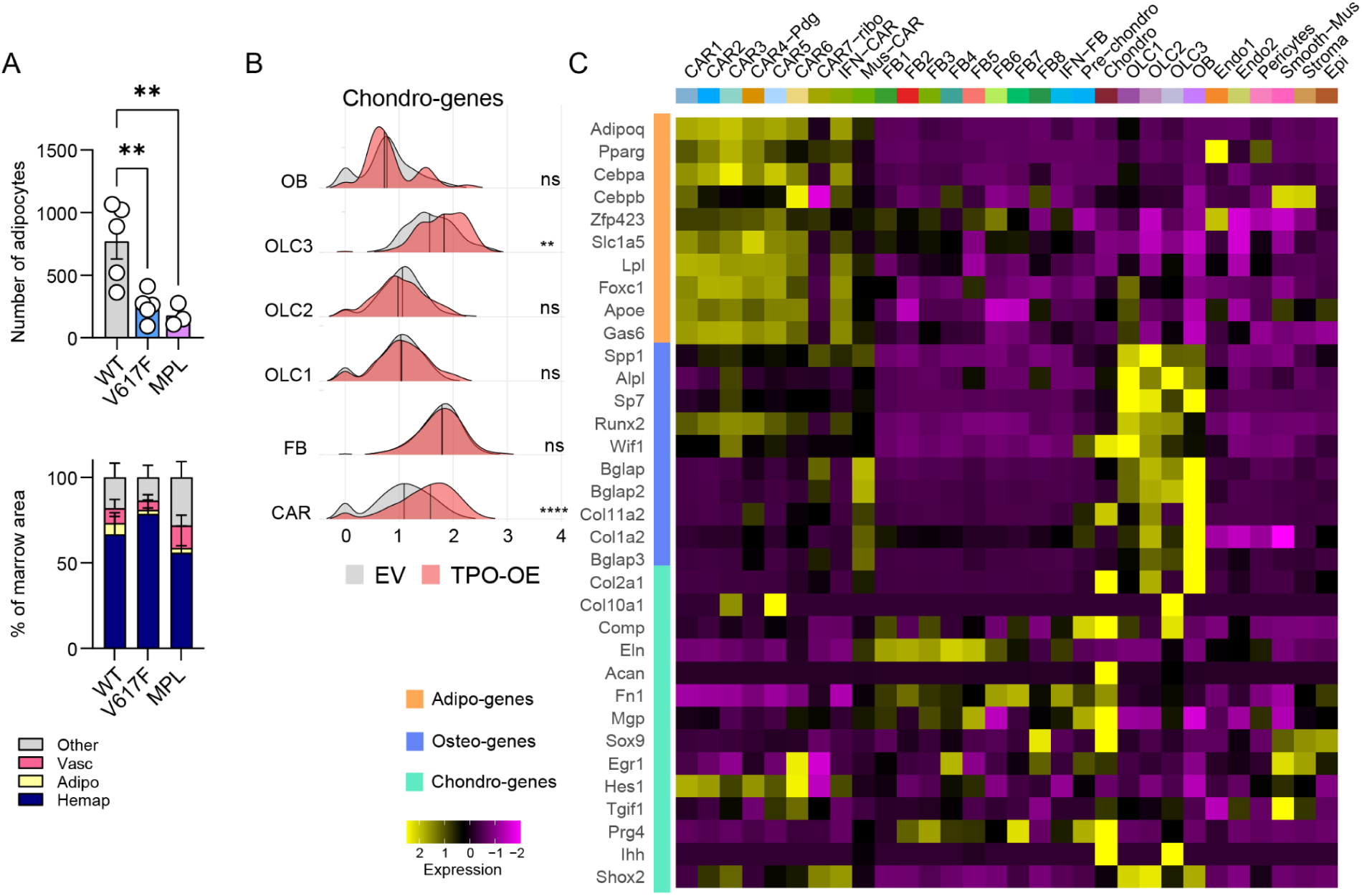
Stromal stem and progenitor cells are skewed in their differentiation in bone marrow fibrosis. (A) “Marrowquant” tissue composition quantification and number of adipocytes in H&E stained WT, JAK2V617F and MPLW515L femurs. One way ANOVA tested. (B) Aggregate expression of chondro-genes per cluster, TPO vs EV, two-sided Wilcox test (C) Heatmap of cluster averages of adipogenesis (adipo), osteogenesis (osteo) and chondro-genes genes of homeostasis subsetted dataset Statistical significance is indicated by: *p<0.05, **p<0.01, ***p<0.001, ****p<0.0001

**Figure S6 (related to figure 6):**
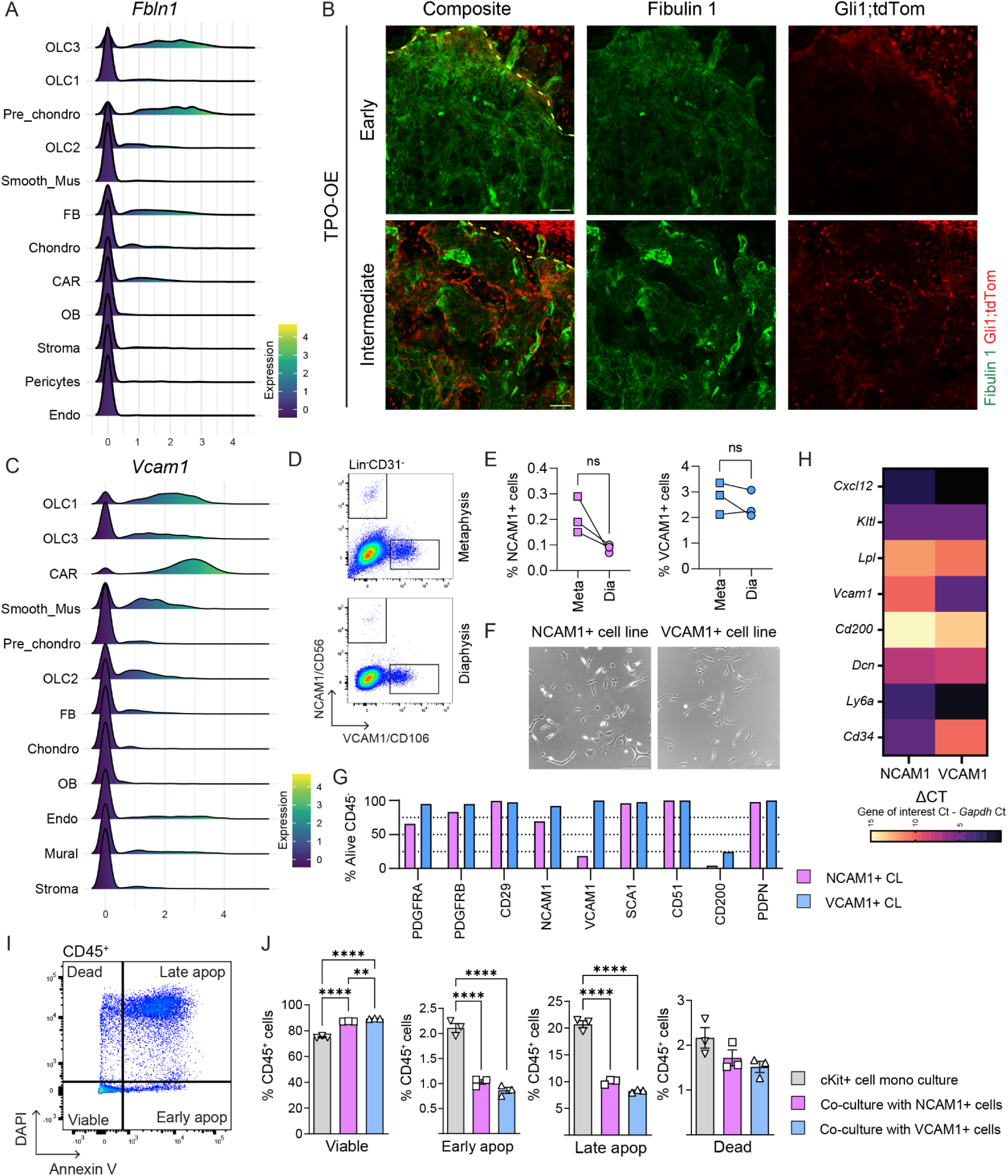
Identification of markers for peritrabecular progenitor cells. (A) Ridge plot of Fibulin1 (*Fbln1*) expression for clusters (B) Confocal images of TPO-OE transplanted tibias in Gli1;tdTom lineage traced mice, at early harvest (5W post transplantation; post-tx) and at intermediate harvest (8W post-Tx) time point. Composite image shown, the right panels show individual channels of Fbln1 and Gli1;tdTom respectively. Scale bar: 50µm. (C) Ridge plot of *Vcam1* expression for clusters. (D) Representative flow cytometry plot showing NCAM1 and VCAM1 expression of lineage depleted BM and digested BC fractions of metaphyseal and diaphyseal fractions of humeri from homeostatic mice (n=3). (E) Quantification of NCAM1+ and VCAM1+ cells found in metaphyseal (meta) and respective diaphyseal (dia) regions of humeri, n=3. Paired t-test. ns:non-significant. (F) Brightfield microscopy of prospectively sort-purified NCAM1+ and VCAM1+ cells respectively, following immortalization, scale bar = 230µm. (G) Percentage of protein marker expression in prospectively sorted NCAM1+ and VCAM1+ cells after culture and immortalization. Proteins as indicated. (H) Heatmap of delta CT of genes of interest shown of prospectively sorted NCAM1+ and VCAM1+ cells after culture and immortalization, at time-point of FACS analysis in panel G. (I) Gating strategy of apoptosis assay (Annexin V combined with DAPI) of CD45+ cells following 24-hour co-culture with NCAM1+ cell line (CL) or VCAM1+ CL. (J) Viability of hematopoietic stem and progenitor cells (cKit-enriched cells) after 24-hour culturing with prospectively sorted NCAM1 or VCAM1 cells, compared to conventional cKit suspension culture. One-way ANOVA. Statistical significance is indicated by: *p<0.05, **p<0.01, ***p<0.001, ****p<0.0001

**Figure S7 (related to figure 7):**
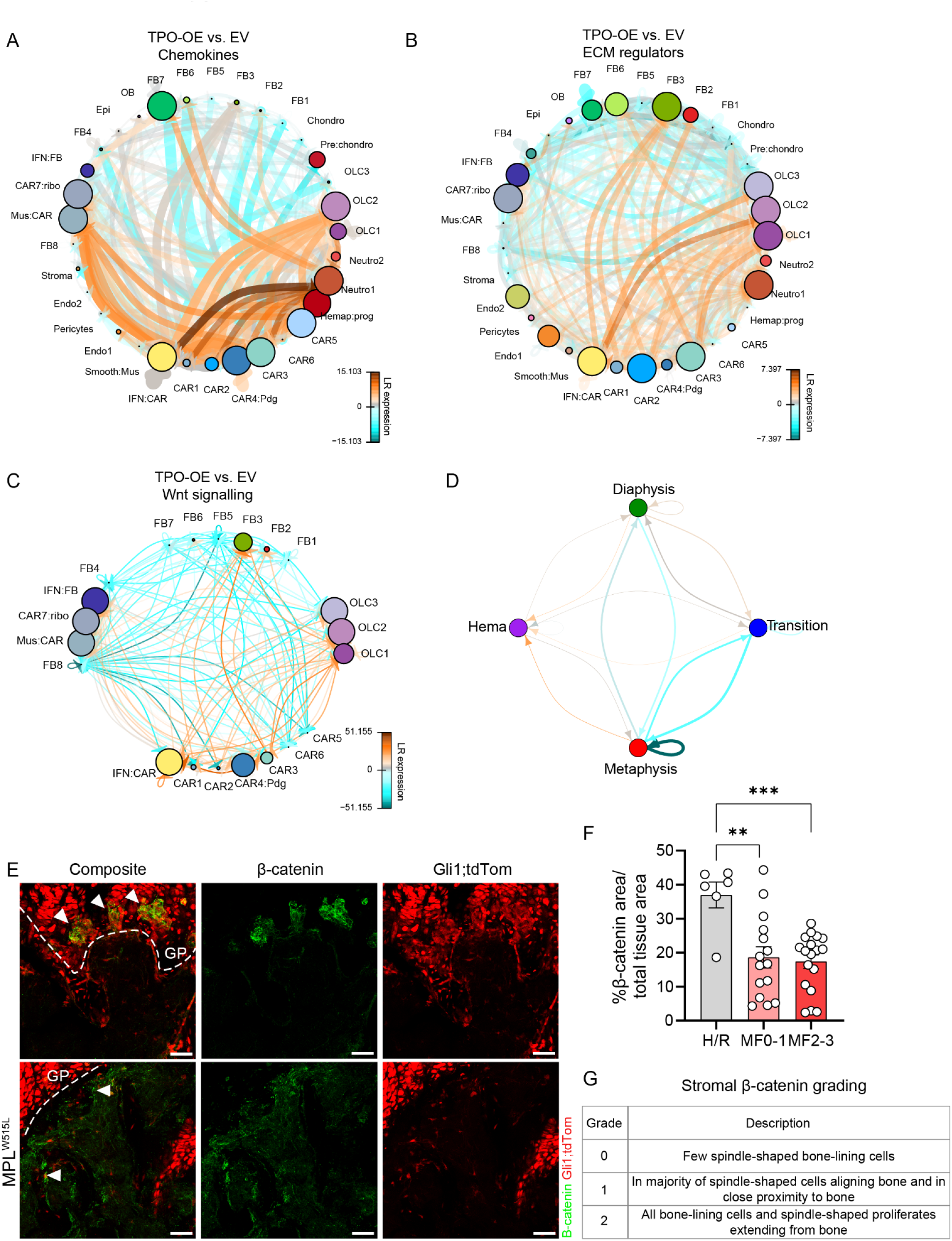
Wnt signaling is reactivated in peritrabecular mesenchymal progenitor cells. (A-C) CCI plot signaling showing receptor interactions of the different niches (metaphysis, diaphysis, transition vs. hematopoietic=hemap) but also cell clusters (TPO-OE vs. EV) filtered on genes belonging to chemokines based on CytoSig database (A) ECM regulators based on Matrissome DB (B) and Wnt signaling based on GO (C) (D) CCI plots of major niches identified in Figure 7, TPO-OE vs. EV (E) Beta-catenin staining of murine humeri, EV= empty vector, control, MPLW515L = fibrotic marrow. Composite image shown, and the right panels show individual channels of beta-Catenin and Gli1;tdTom respectively. White arrowheads highlight beta-catenin and Gli1;tdTom co-expression.Scale bar = 50µm (F) Whole tissue quantification of active beta-catenin signal, shown per MF grade, One-way ANOVA. (G) Grading table for stromal beta catenin grading (sbCG) of human bone marrow biopsies. One-way ANOVA with Kruskal-Wallis. H/R = healthy/reactive, MF = myelofibrosis grade 0-3 Statistical significance is indicated by: *p<0.05, **p<0.01, ***p<0.001, ****p<0.0001

**Figure S8 (related to figure 8):**
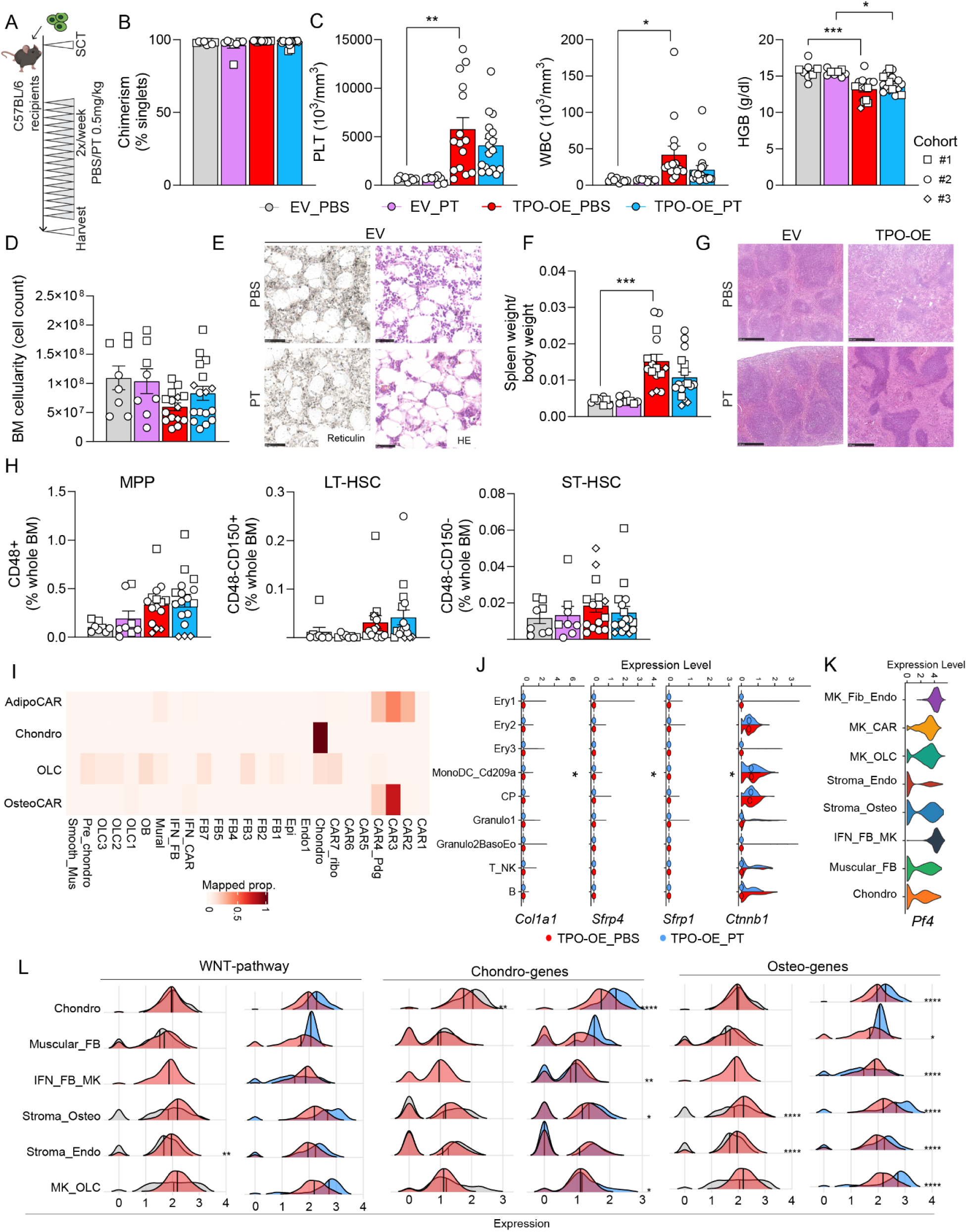
Wnt signaling is reactivated in peritrabecular mesenchymal progenitor cells. (A) Schematic of experimental design, following BM transplant of cKit-enriched cells transduced with EV-eGFP or TPO-OE-eGFP viral vector, mice were treated twice weekly with Pyrvinium-Tosylate (PT)-DMSO solution or PBS-DMSO as a control starting 21 days after transplantation. (B) BM chimerism of recipient mice at harvest time-points, three independent cohorts shown (C) Progression of MPN blood phenotype of cohort in F7E, PLT = platelets, WBC = white blood cells, HGB = hemoglobin. One-way ANOVA performed on last time-point, only significant results shown. (D) BM cellularity per group, and spleen weight/ body weight. One-way ANOVA performed. (E) Representative images of reticulin and H&E staining of paraffin-embedded bones in EV control mice (F) Spleen over body weight at harvest of experimental groups. One-way ANOVA performed. (G) Representative images of H&E images of spleens, scale bar: 500µm (H) HSC FACS of cohorts, individual mice per cohorts shown. One-way ANOVA performed. Non-significant differences. (I) Proportion of cell clusters from TPO-OE_PBS and TPO-OE_PT dataset (from figure 8C) associated with tdTom-lineage cell clusters (from figure 1B) (J) Remainder of violin plots of figure 8D, median depicted with circle, wilcox test for significance. (K) Violin plot of *Pf4* expression in SpDs. MK (megakaryocyte); Fib (fibroblast); CAR-cells; OLC (osteolineage cells); IFN (interferon); endo (endothelium) (L) Ridge plots of remaining SpDs of figure 8J, K, L. TPO-OE PBS vs EV PBS in the left panel, TPO-OE PT vs TPO-OE PT in the right panel. One sided Wilcox test performed per spatial domain cluster, per comparison. Statistical significance is indicated by: *p<0.05, **p<0.01, ***p<0.001, ****p<0.0001

